# METTL9 sustains vertebrate neural development primarily via non-catalytic functions

**DOI:** 10.1101/2025.05.05.652220

**Authors:** Azzurra Codino, Luca Spagnoletti, Claudia Olobardi, Alessandro Cuomo, Helena Santos-Rosa, Martina Palomba, Natasha Margaroli, Stefania Girotto, Rita Scarpelli, Shi-Lu Luan, Eleonora Crocco, Paolo Bianchini, Andrew J. Bannister, Stefano Gustincich, Tony Kouzarides, Riccardo Rizzo, Isaia Barbieri, Federico Cremisi, Robert Vignali, Luca Pandolfini

**Affiliations:** Italian Institute of Technology (IIT), Center for Human Technologies, Via Enrico Melen 83, 16152 Genoa, IT; Department of Biology, University of Pisa, via Luca Ghini 13, 56126 Pisa, IT; Department of Experimental Oncology, IEO, European Institute of Oncology IRCCS, Milan, 20139, IT; The Wellcome Trust/Cancer Research UK Gurdon Institute and Department of Pathology, University of Cambridge, Tennis Court Road, Cambridge, CB2 1QN, UK; Italian Institute of Technology (IIT), Center for Convergent Technologies, Via Morego 30, 16163 Genoa, IT; University of Cambridge, Dep. of Pathology, 10 Tennis Court Road, Cambridge CB2 1QP, UK; BIO@SNS, Scuola Normale Superiore di Pisa, Via Giuseppe Moruzzi, 56124 Pisa, IT; NIC@IIT, Italian Institute of Technology (IIT), Via Enrico Melen 83, 16152 Genoa, IT; Institute of Nanotechnology, National Research Council (CNR-NANOTEC), Campus Ecotekne, Via Monteroni, 73100, Lecce, Italy; Department of molecular biotechnology and health sciences, Molecular Biotechnology Center, University of Turin, Via Nizza 52, 10126, Turin, Italy

## Abstract

METTL9 is an enzyme catalysing N1-methylation of histidine residues (1MH) within eukaryotic proteins. Given its high expression in vertebrate nervous system and its potential association with neurodevelopmental delay, we dissected *Mettl9* role during neural development. We generated three distinct mouse embryonic stem cell lines: a complete *Mettl9* knock-out (KO), an inducible METTL9 Degron and a line endogenously expressing a catalytically inactive protein, and assessed their ability to undergo neural differentiation. In parallel, we down-regulated *mettl9* in *Xenopus laevis* embryos and characterised their neural development. Our multi-omics data indicate that METTL9 exerts a conserved role in sustaining vertebrate neurogenesis. This is largely independent of its catalytic activity and occurs through modulation of the secretory pathway. METTL9 interacts with key regulators of cellular transport, endocytosis and Golgi integrity; moreover, in *Mettl9^KO^* cells Golgi becomes fragmented. Overall, we discovered the first developmental function of *Mettl9* and linked it to a 1MH-independent pathway, namely, the maintenance of the secretory system, which is essential throughout neural development.

## Introduction

Methyltransferase-like (METTL) proteins constitute a diverse class of enzymes sharing the evolutionary conserved *S*-adenosylmethionine (SAM) binding domain, which endows them with catalytic activity^1,2^. Upon SAM cofactor binding, METTL proteins can transfer methyl groups from SAM to different macromolecules, such as RNA or proteins.

Amplification and deletion of various METTL genes have been associated with different pathological conditions, including cancer, where the same METTLs can act either as tumour suppressors or as oncogenes, depending on the tissue and cellular context^3^. Besides their established role in cancer, METTLs sustain many physiological processes, including cell fate specification and differentiation^3^. METTL3 maintains hematopoietic stem cell quiescence^4^ besides acting as an oncogene in acute myeloid leukemia^5^. METTL1 safeguards self-renewal potential in mouse embryonic stem cells (mESCs) and regulates ectodermal and neural differentiation of mESCs^6^. Similarly, METTL6 is important for preserving mESCs pluripotency^7^ and METTL17 knock-out (KO) delays embryonic stem cell differentiation by decreasing mitochondrial respiration^8^; METTL11A sustains myoblasts differentiation^9^ and prevents premature ageing through the maintenance of the quiescent neural stem cell pool^10^.

Histidine (His) methylation is a post-translational modification whose biological significance has remained elusive for a long time^11^. Only 2 His-specific methyltransferases have been identified: METTL18, which methylates the nitrogen in position 3 of histidine imidazole ring (generating 3-methylhistidine, 3MH) mainly in RPL3 protein, modulating translational elongation^12,13^ and METTL9, which was found to methylate the nitrogen 1 of histidines (generating 1MH) in hundreds of proteins containing the H[ANGST]H motif^14^. More recently, the structural details of METTL9 substrate recognition and catalysis have also been clarified^15,16^. METTL9 targets include mitochondrial respiration factors like NUFB3 and zinc transporters, whose zinc binding affinity is changed upon METTL9 KO^14^.

Similarly to other METTL proteins, METTL9 has also been linked to cancer biology. It is upregulated in many tumours (where its expression level correlates with poor prognosis) and decreasing METTL9 expression in hepatocellular carcinoma cells slows down their proliferation and ability to generate tumours *in vivo* as well as their invasiveness^17^. Moreover, METTL9-dependent methylation of certain zinc transporters is associated with tumour growth^18^. Interestingly, methylation of the immunomodulatory S100A9 protein^14,19^ is dynamically regulated in neutrophils upon infection^20^. Low levels of S100A9 methylation endow mice with a better anti-bacterial ability by increasing S100A9 affinity to zinc^20^. However, besides its role in cancer and in immune response, the biological function of METTL9 in physiological and developmental processes has remained unknown. Here we show that METTL9 is highly expressed in the vertebrate nervous system where it sustains early neurogenesis. By using complementary genetic systems, we also reveal that this function is mainly independent of its catalytic activity and that it is linked to the secretory pathway and Golgi integrity.

## Results

### Mettl9 is important for vertebrate neural development

To uncover the physiological relevance of Mettl9, we first sought to assess whether the human *METTL9* locus was associated with any disease. Importantly, by mining the DECIPHER database^21^, we identified 6 patients harbouring heterozygous deletions of a genomic region encompassing the *METTL9* gene in chromosome 16 (Fig. 1a). Interestingly, 5 out of the 6 patients displayed at least one nervous system-related phenotype such as cognitive impairment, autistic behaviour, morphological central nervous system abnormality and intellectual disability, while the sixth patient displayed global developmental delay (Fig. 1a). Some of them also showed varying degrees of hearing impairment, possibly ascribed to the loss of the inner ear specific gene Otoancorin, *OTOA*, which is also present within the deleted region together with the *IGSF6* gene, and was previously linked to deafness^22,23^ (See Fig. 1a legend).

**Fig. 1:**
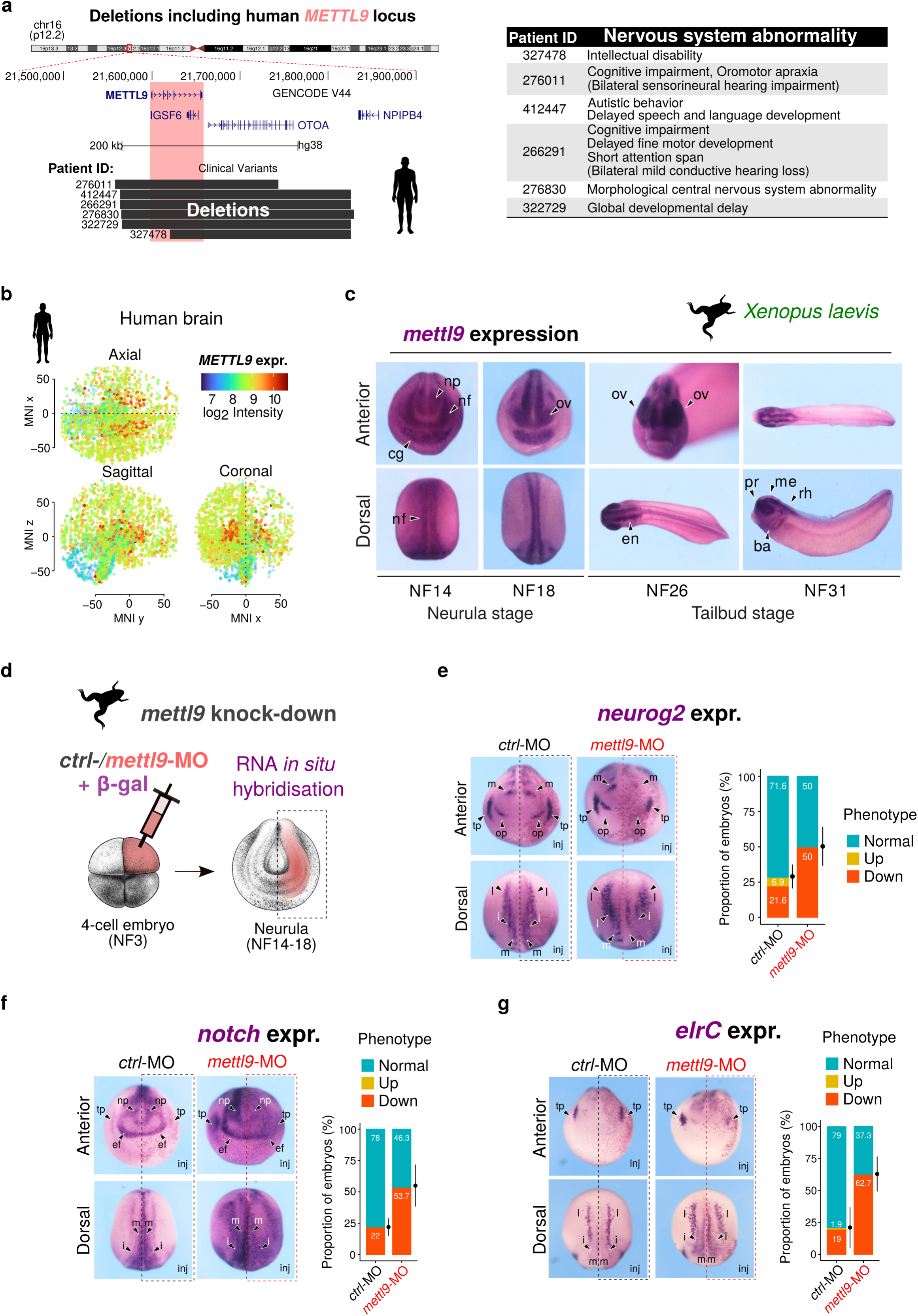
Mettl9 is expressed in vertebrate nervous system and is important for *X. laevis* neural development *in vivo*. **a** Deletions encompassing human *METTL9* locus on Chromosome 16 in patients, who display nervous system-related phenotypes (table on the right) (from DECIPHER). Deletions also include the *OTOA* gene, the *IGSF6* gene encoding a predicted transmembrane receptor of the immune system and, in few cases, also the *RRN3P1* pseudogene. **b** *METTL9* normalised microarray expression level (log_2_ intensity) in the three sections of a three-dimensional map of adult human brain (Allen Human Brain Atlas). Coordinates are referred to the Montreal Neurological Institute (MNI) standard spatial template. **c** Representative anterior and dorsal views images of *X. laevis* embryos showing *mettl9* mRNA expression (purple) by whole-mount RNA *in situ* hybridisation (WISH). At Niewkoop-Faber stage 14 (NF14), black arrowheads indicate: neural plate (np), neural fold (nf) and cement gland (cg) and at NF18 they show optical vesicles (ov). At tailbud stage (NF26) it is shown the encephalon (en) which, at NF31, is subdivided into prosencephalon (pr), mesencephalon (me) and rhombencenphalon (rh). Branchial arches (ba) are also shown at NF31. **d** Schematic of *mett9* knock down (k.d.) strategy at 4-cell stage embryo: microinjection of *mettl9* morpholino oligonucleotide (MO) (pink) in the dorsal left blastomere is shown. WISH is then performed at later stages (neurula) to assess potential developmental abnormalities in the injected side of embryos (right side, pink). Embryo images adapted from Zahn et al.^171^. **e-g** Representative images of *ctrl*-MO and *mett9*-MO k.d. embryos at neurula stage (NF14), showing the expression of neural markers *neurog2* (*ngn*) (**e**), *notch* (**f**) and *elrC* (**g**) by WISH. Lateral *(l)*, intermediate *(i)* and medial *(m)* stria, trigeminal and olfactory placodes (tp, op), neural plate (np) and eye field (ef) are shown. Arrowheads indicate the regions affected in *mettl9*-MO. Inj. is Injected side. Bar graphs on the right show the quantification of *ctrl*-or *mettl9*-MO k.d. embryos screened for altered *neurog2* (n=102 for *ctrl-*MO and n=62 for *mettl9*-MO respectively), *notch* (n=59 *ctrl*-MO; n=54 *mettl9-*MO) or *elrC* (n=105 *ctrl*-MO; n=59 *mettl9-*MO) expression (χ2 test).

Since the phenotypes associated with these *METTL9*-containing deletions suggested a putative role for *METTL9* in neurodevelopmental disorders, we assessed *METTL9* expression in the developing brain during prenatal life. By interrogating the BrainSpan Atlas of the Developing Human Brain^24^, we found that *METTL9* is expressed at high levels during human neural development, while its expression decreases in postnatal brain (Supplementary Fig. 1a). Within the adult brain *METTL9* levels peak in the striatum (Supplementary Fig. 1a, 30-40 yrs), which was also confirmed by the three-dimensional map of adult human brain (Allen Human Brain atlas^25^) (Fig. 1b) and exploration of the GTEx datasets^26^ (Supplementary Fig. 1b). Furthermore, we confirmed that *Mettl9* is highly expressed in the direct and indirect spiny projection neurons of the mouse striatum, as shown by re-analysis of available scRNA-seq data^27^ (Supplementary Fig. 1c). In summary, given the high expression levels of METTL9 in mouse and human brain, as well as the neurodevelopmental phenotypes potentially related to its deletion in patients, we hypothesised that Mettl9 dosage could be important for sustaining early neural development.

Since the most crucial and earliest events of mammalian brain development are shared among vertebrates^28^, the African clawed frog *Xenopus laevis* represents an exceptional model organism for phenotypic screening of highly conserved genes with a putative developmental function, mainly due to the ease of *in vivo* embryo manipulation^29^.

The *Mettl9* gene is evolutionarily conserved across the animal kingdom^2^, and the encoded protein is highly conserved between *Mammalia* and *Amphibia* (*Xenopus laevis)* both in terms of amino acid sequence (which shows an overall similarity of 82.4% and 82.7% with *Mus musculus* and *Homo sapiens*, respectively; Supplementary Fig. 2a) and structure (Supplementary Fig. 2b), particularly in the region encoding the catalytic SAM binding domain (Supplementary Fig. 2a,b). Therefore, we reasoned that its core developmental functions likely originated early in evolution and have since been conserved. Thus, we performed whole-mount RNA *in situ* hybridization (WISH) and analysed the spatio-temporal expression pattern of *mettl9* throughout *X. laevis* early development (Supplementary Fig. 2c). Interestingly, *mettl9* mRNA was detected very early during embryogenesis, in the NF6.5-NF7 blastula (Supplementary Fig.2 d), probably due to maternal genome contribution. It remained highly expressed at gastrula stage (NF10.5), following zygotic gene activation and germ cell layer and axis specification. Importantly, *mettl9* was highly expressed during early neurulation (NF14-18 stages) (Fig. 1c), both in the neural fold (nf) and neural plate (np), before neural tube folding and closure (NF14 and 18). After the completion of neurulation, *mettl9* expression becomes more restricted to the anterior part of the early (NF20-23) (Supplementary Fig. 2d) and late (NF26 and 31) tailbud stages (Fig. 1c), in the encephalon (en) and optical vesicles (ov; NF26) (Fig. 1c), and also in neural crest cells (ncc; NF31) (Supplementary Fig. 2e).

These data strongly indicate that *mettl9* is highly expressed in the nervous system during early amphibian development and its levels in the brain remain significant at later stages. Importantly, these data are consistent with mammalian expression patterns, suggesting that Mettl9 expression in the developing nervous system is a conserved feature of vertebrates.

We then set out to investigate the requirement of *mettl9* for neurogenesis *in vivo*, by analysing *mettl9* knock-down (k.d.) embryos. To this end, we designed a *mettl9*-morpholino antisense oligonucleotide (MO) which targets the exon1/intron1 junction of the *mettl9* pre-mRNA, thus impairing *mettl9* splicing (Supplementary Fig. 2f), as confirmed by qPCR (Supplementary Fig. 2g). We injected *mettl9*-MO or *ctrl*-MO into one of the two dorsal blastomeres of 4-cell stage embryos (Fig.1d), leaving the contralateral side of the embryo as an internal control. To assess whether *mettl9*-MO embryos were able to undergo early neurogenesis, we analysed the expression pattern of key neural markers by WISH: strikingly, we found that *neurog2*, a marker of neuronal precursors^30^, *notch,* involved in regulation of lateral inhibition within the developing neural plate^31^ and *elrC,* a marker of committed neuronal progenitors^32^, were down-regulated at the neurula stage (Fig. 1e,f,g). In particular, *neurog2* was down-regulated in the medial, intermediate and lateral columns of prospective neuronal precursors, and also in the olfactory and trigeminal placodes (Fig. 1E, arrowheads); *notch* pattern was severely affected on the injected side both in the anterior domain around the presumptive eye field and in the trunk region (Fig. 1f, arrowheads); and *elrC* was also affected in the developing neurons (Fig. 1g), as shown by alterations in the intermediate neuron precursors and trigeminal precursors (arrowheads). These effects were found with a significantly higher proportion in *mettl9*-MO injected embryos than in *ctrl*-MO injected embryos (Fig. 1e-g).

Overall, these data supported the idea that *mettl9* may exert a role in early neurogenesis and prompted us to further dissect its function in the context of a mammalian model.

### Constitutive depletion of Mettl9 from mESCs impairs neural fate commitment and neural differentiation

Given that *mettl9* depletion negatively affected early neural development in *X. laevis* embryos, we speculated that loss of Mettl9 would similarly impair mammalian neural fate specification. To address this, we used mouse embryonic stem cells (mESCs) to study neural commitment and differentiation. mESCs can faithfully recapitulate the main steps of mammalian neurogenesis in a few days *in vitro* and allow fine genetic and molecular manipulations. Thus, we steered mESCs towards a neural fate by culturing them in the absence of serum/leukaemia inhibitory factor (LIF) and by inhibiting both BMP and Wnt signalling, as previously described^33,34^ (Fig. 2a). After 5 days in culture with Wnt/BMP inhibitors (DIV5), mESCs acquire neural stem cell (NSC) identity, as shown by the expression of the neural markers *Nestin*, *Dlx2* and *Gsx2* (Supplementary Fig. 3a). Then, after supplying Sonic Hedgehog (SHH) agonist from DIV5 until DIV10, NSCs acquire the identity of ventral telencephalic neural progenitor cells (NPCs), as shown by the progressive up-regulation of the striatal genes *Drd1a*, *Drd2* and *Gad1* (Supplementary Fig. 3a). Importantly, we observed that *Mettl9* was also up-regulated both at the mRNA (Fig. 2b) and protein level (Fig. 2c) upon mESCs commitment to NSCs, and it reached even higher levels in NPCs. Overall, these data are consistent with our previous findings in the *X. laevis* and mammalian studies, reinforcing the notion that our cell model is suitable to study the physiological roles of *Mettl9* during mammalian neurogenesis.

**Fig. 2:**
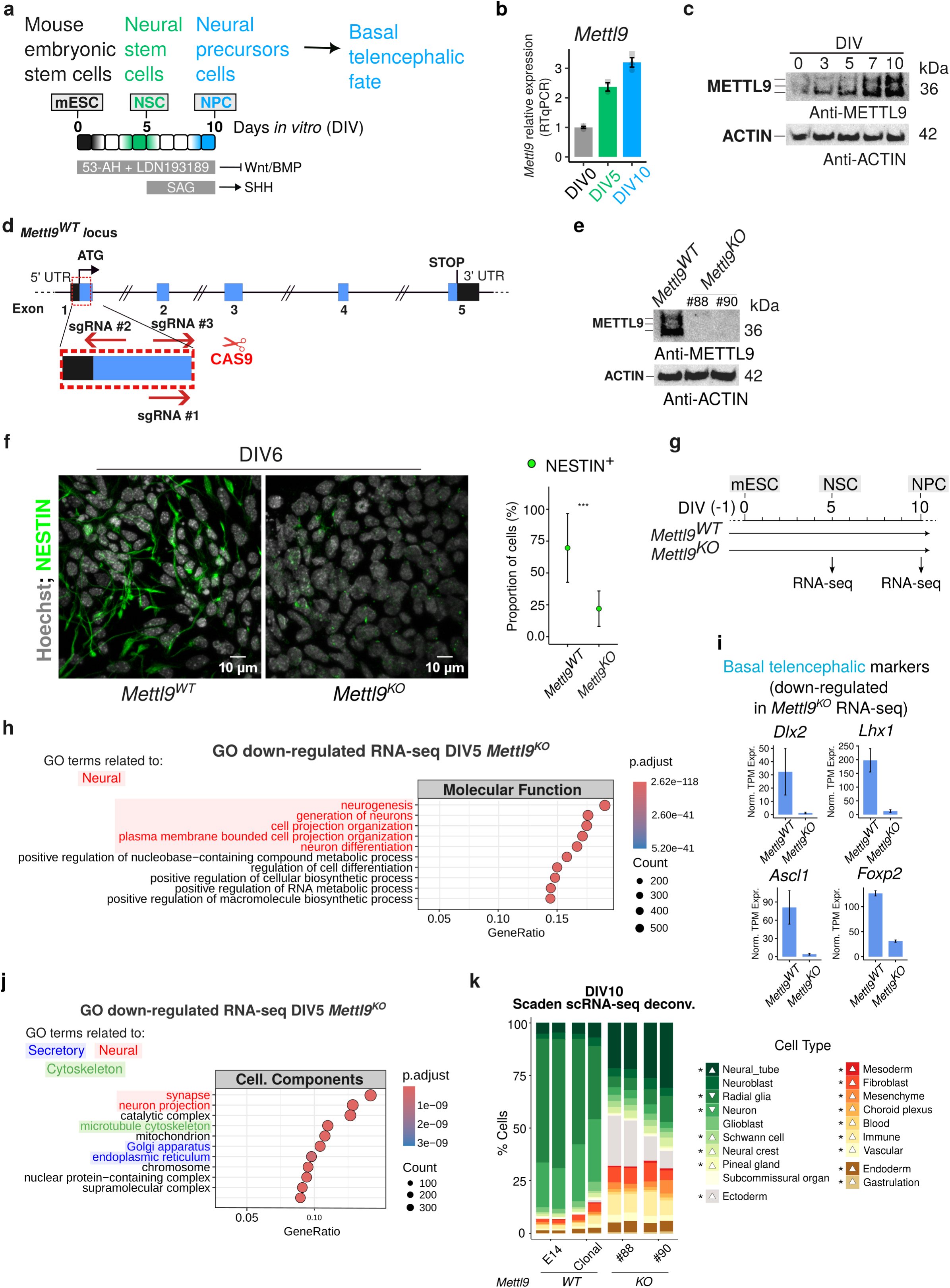
Constitutive *Mettl9* depletion impairs mESCs neural priming and differentiation. **a** Schematic depicting the neural differential protocol adopted in this study. **b** *Mettl9* mRNA relative expression normalised on *β-actin* (by qPCR), at DIV0, DIV5, DIV10. **c** METTL9 protein expression shown by Western Blot (WB) from DIV0 to DIV10 (anti-METTL9 antibody; kDa: kDalton). **d** Schematic of the mouse *Mettl9* genomic locus and genetic strategy to generate *Mettl9^KO^* mESCs by CRISPR/Cas9. In red is shown the portion of Exon1 targeted by sgRNAs to generate a deletion. **e** WB from neural stem cells extracts (DIV7) showing endogenous METTL9 expression with an anti-METTL9 antibody (lines: WT; KO: #88 and #90). **f** Representative immunofluorescence (IF) images of *Mettl9^WT^*and *Mettl9^KO^* NSCs (DIV6), stained with an anti-NESTIN antibody (green) and Hoechst (grey). Scale bar is 10 μm; relative quantification of NESTIN signal on the right (Number of cells counted: 2475 for *Mettl9^WT^*; 2496 for *Mettl9^KO^*. Wilcoxon test). **g** Schematic of the key timepoints during neural differentiation of *Mettl9^KO^* mESC (DIV5 and DIV10) analysed by RNA-seq. **h** Top (10) GO Molecular function terms down-regulated in *Mettl9^KO^* RNA-seq (DIV5). In red, neural-related terms. **i** Normalised transcripts per million (TPM) expression of basal telencephalic markers *Dlx2*, *Lhx1*, *Ascl1*, and *Foxp2*. **j** Top (10) GO Cellular Component terms down-regulated in *Mettl9^KO^* RNA-seq (DIV5). **k** Cell type representation in control WT (E14 or clonal) and *Mettl9^KO^* (#88 and #90) lines inferred by SCADEN deconvolution analysis of scRNA-seq data (DIV10). Each result is shown in duplicate.

We then constitutively depleted *Mettl9* from mESCs and assessed their ability to undergo neural priming and differentiation. To this end, we engineered a *Mettl9* knock-out (*Mettl9^KO^*) mESC line using CRISPR/Cas9 technology, by targeting the first exon of the endogenous *Mettl9* locus with 3 sgRNAs (Fig. 2d). We obtained two *Mettl9^KO^*clones (#88 and #90), each harbouring biallelic deletions within the first exon of *Mettl9* (Supplementary Fig. 3b,c), which resulted in the complete loss of METTL9 protein, as shown by WB (Fig. 2e). Consistently, *Mettl9* mRNA levels were almost undetectable by qPCR in *Mettl9^KO^* cells (Supplementary Fig. 3d), suggesting that METTL9 degradation was most likely due to the non-sense mediated mRNA decay pathway^35^. Importantly, the sgRNAs used did not generate off-target mutations across the genome, as confirmed by whole-genome sequencing (Supplementary Fig. 3e).

We first assessed whether *Mettl9^KO^* mESCs could differentiate into *bona fide* NSCs; strikingly, we found a significantly lower number of neural NESTIN-positive (NES^+^) cells in *Mettl9^KO^*cultures compared to the *Mettl9^WT^* (parental line, E14), as shown by immunofluorescent (IF) staining of NSCs and cell counting, at DIV6 (Fig. 2f).

To gain a comprehensive understanding of the molecular processes affected by *Mettl9* loss, we profiled gene-expression of *Mettl9^KO^* NSCs by RNA-seq (Fig. 2g). Importantly, at DIV5 we found severe transcriptomic alterations, as we identified 5732 mis-regulated genes in both the #88 and #90 clonal *Mettl9^KO^* NSCs compared to the controls clonal and parental (E14) *Mettl9^WT^* cell lines, with 2862 down-regulated and 2870 up-regulated genes (Supplementary Fig. 4a; Supplementary Data 1). Notably, both *Mettl9^KO^* clones exhibited a very consistent gene regulation, distinct from both parental (E14) and clonal (i.e. non-edited) wild-type (*Mettl9^WT^*) lines (Supplementary Fig. 4b). As anticipated by the decrease in NES^+^ neural cells shown by IF, among the most down-regulated Gene Ontology (GO) Molecular Function terms we found many neural-related entries such as “neurogenesis” and “neuron differentiation” (Fig. 2h) (refer to Supplementary Data 2 for full GO list). These were exemplified by down-regulation of the NSC markers *Sox1*, *Nestin*, *Foxg1* and *Shh* (Supplementary Fig. 4c) as well as of the early basal telencephalic gene markers *Dlx2*, *Lhx1*, *Ascl1* and *Foxp2* (Fig. 2i) and *Dlx1*, *Lhx5* (Supplementary Fig. 4c), which was accompanied by the up-regulation of stemness (mESC) markers such as *Pou5f1*, *Nanog, Nodal* (Supplementary Fig. 4c). Interestingly, “cell projection organization” was also found among the most down-regulated GO terms. Consistently, some of the most down-regulated GO terms concerning Cellular Component were: “synapses”, “neuron projection”, “microtubule cytoskeleton”, “Golgi apparatus” and “ER” (Fig. 2j; Supplementary Data 2). Moreover, among the up-regulated Molecular Functions we found many metabolic and biosynthetic processes (Supplementary Fig. 4d; Supplementary Data 2). Overall, these transcriptomic data indicate that upon *Mettl9* loss, differentiating mESCs cannot completely shut down their stemness program and cannot undergo a neurogenetic route, thus precluding their acquisition of a typical NSCs identity.

After discovering the impairment in NSC commitment at DIV5, we wondered whether *Mettl9^KO^* NSCs could undergo neural differentiation. By DIV10, *Mettl9^KO^* NPCs became morphologically different from their *Mettl9^WT^* counterparts, as they lacked neural projections and resembled fibroblast-like cells (Supplementary Fig. 4e). This strong phenotype was confirmed by the large number of mis-regulated genes found in both clonal *Mettl9^KO^* lines at DIV10 (3298 down-regulated genes and 3232 up-regulated genes) compared to control *Mettl9^WT^* NPCs (Supplementary Fig. 4f,g; Supplementary Data 1). Next, we inferred the identity of these aberrant *Mettl9^KO^* cells, through the deconvolution of cell type composition using neural-networks^36^ trained over single-cell atlas of developing mouse brain^37^. Importantly, this analysis revealed that *Mettl9^KO^* cells at DIV10 had a relatively higher proportion of neural tube cells (Fig. 2k), usually present at earlier stages of neural development, a decrease in radial glial cells and a massive increase in other cell lineages such as mesoderm and ectoderm (e.g. fibroblast-related), as anticipated from the macroscopical observation of the cellular phenotype. Overall, constitutive loss of *Mettl9* prevents mESCs from generating *bona fide* NSCs and, at later stages, massively impairs the specification of NPCs, while promoting the aberrant acquisition of mesodermal and non-neural ectodermal identity.

### Acute depletion of METTL9 from mESCs via Degron only partially mimics METTL9 constitutive depletion

To shed light on the biological and molecular processes more directly controlled by METTL9 during neural differentiation, we employed an inducible protein-Degron system^38,39^. We fused METTL9 to the mutated prolyl isomerase FKBP12^F36V^, to induce its rapid degradation in live cells (Fig. 3a)^38,39^. Using CRISPR/Cas9, we generated an endogenously tagged *Mettl9^FKBP^*^12^*^-F36V-FLAG^* mESCs line (in short *Mettl9^Deg^,* which stands for *Mettl9 Degron*), by targeting the last exon of the *Mettl9* locus (Fig. 3b and Supplementary Fig. 5a) before the STOP codon. The resulting METTL9-FKBP12^F36V^-FLAG (METTL9-DEG) protein was expressed at the expected size in mESCs, as shown by WB (Fig. 3c). Upon cellular uptake, the dTAG^V^-1 drug specifically recognises the mutated FKBP12^F36V^ (and not the endogenous, ubiquitous FKBP12) and recruits the E3 ubiquitin ligase CRBN to FKBP^F36V^, which in turns ubiquitylates METTL9-FKBP^F36V^ thereby promoting proteasomal degradation of the fusion protein (Fig. 3a).

**Fig. 3:**
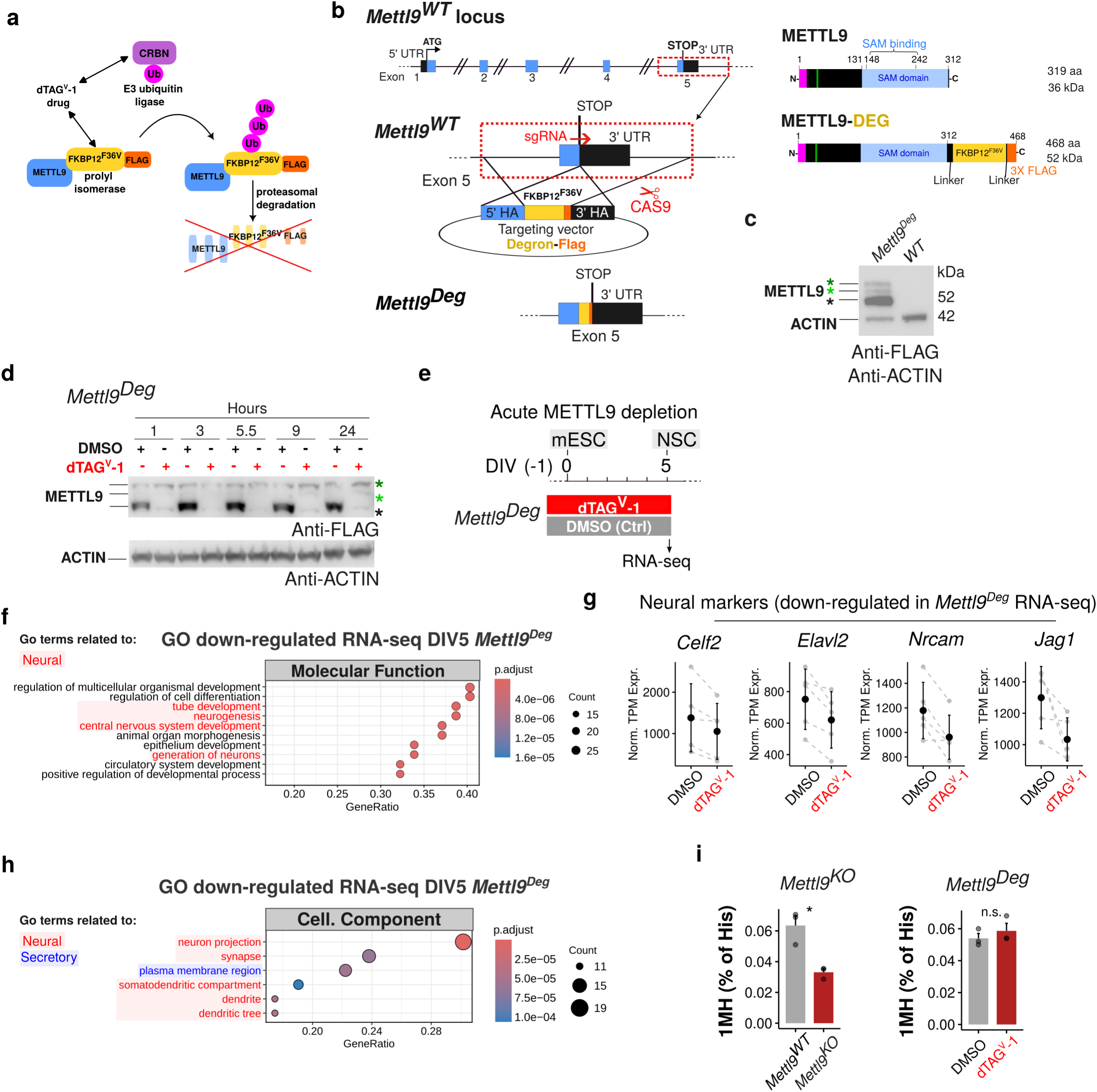
Acute depletion of METTL9-DEGRON affects the gene expression profile of mNSCs. **a** Schematic depicting METTL9-DEGRON system and relative mechanism of degradation upon dTAG^V^-1 supply. **b** *Mettl9* genomic locus and its genetic targeting via CRISPR/Cas9 for *Mettl9^Deg^* generation. On the right, the resulting C-terminally tagged METTL9-DEG protein is shown (aa: amino acid; kDa: kDalton). **c** Validation of METTL-DEGRON protein expression in *Mettl9^Degron^* mESCs by WB (anti-FLAG and anti-ACTIN antibodies). Black, green and dark green asterisks (*) refer to the lowest, intermediate and top METTL9 bands, respectively. **d** Time course expression analysis of METTL9-DEG by WB, after supplying dTAG^V^-1 (or DMSO, Ctrl) to mESCs for 1, 3, 5.5, 9 or 24 hours. **e** Schematic of experimental strategy for acute METTL9-DEG depletion (by dTAG^V^-1) in *Mettl9^Deg^* mESCs and molecular analysis at DIV5. **f** Top 10 Molecular function GO terms down-regulated in *Mettl9^Deg^* RNA-seq (DIV5). **g** Normalised TPM expression of neural marker genes, from *Mettl9^Deg^*RNA-seq (DIV5). **h** Top down-regulated Cellular Component GO terms in *Mettl9^Deg^*. **i** Relative bulk 1MH levels (% of total histidine) in *Mettl9^WT^* and *Mettl9^KO^* (left); DMSO– and dTAG^V^-1-treated *Mettl9^Deg^* (right), NSCs (DIV6), quantified by mass spectrometry. (Wilcoxon test).

Importantly, within 1 hour from dTAG^V^-1 administration, most of METTL9-FKBP-FLAG was degraded in mESCs, as shown by WB (Fig. 3d).

Therefore, we first assessed whether induced METTL9 protein depletion negatively impacted neural cell fate specification. To this end, we continuously supplied the dTAG^V^-1 drug, or DMSO for control, from DIV(–1) up until DIV5 (Fig. 3e) and then analysed the transcriptome of treated homozygous *Mettl9^Deg^* NSCs, by RNA-seq. At DIV5, we found 63 genes down-regulated and 126 up-regulated (Supplementary Fig. 5b; Supplementary Data 1) in the dTAG^V^-1-treated NSCs compared to the DMSO controls. GO analysis of the differentially regulated genes revealed that in the dTAG^V^-1-treated NSCs “tube development”, “neurogenesis”, “nervous system development” and “generation of neurons” were among the most down-regulated Molecular Function terms in comparison with DMSO controls (Fig. 3f; Supplementary Data 2), as exemplified by the neural markers *Celf2*, *Elavl2*, *Nrcam*, *Jag1* (Fig. 3g; Supplementary Data 1) as well as *Fzd1*, *Map2*, *Robo2*, *Zic1* and *Dll1* (Supplementary Fig. 5c). Moreover, among the most down-regulated GO terms concerning Cellular Component we found “neuron projection”, “synapse”, “plasma membrane region”, “somatodendritic compartment”, “dendrite” and “dendritic tree” (Fig. 3h; Supplementary Data 2), which were all down-regulated also in the *Mettl9^KO^* NSCs. Furthermore, among the up-regulated Molecular Functions (GO terms) we found many metabolic and biosynthetic processes (Supplementary Fig. 5d; Supplementary Data 2), similarly to *Mettl9^KO^* NSCs, and the up-regulated Cellular Component terms included “synapses” and “post-synapses” (Supplementary Fig. 5e). Therefore, the mis-regulation (mainly down-regulation) of neural-related genes as well as the up-regulation of metabolic-related genes in the *Mettl9^KO^*NSCs at DIV5 very likely represented *bona fide* specific effects of METTL9 depletion. Despite the similarity in the biological processes involved, the effects in the *Mettl9^Deg^*were very mild when compared with the *Mettl9^KO^*-transcriptomic analysis, both in terms of number of affected genes and in the strength of their mis-regulation. Consistent with this, no major macroscopic defects were observed upon neuralisation, as shown by a comparable number of Nestin positive cells between DMSO-or dTAG^V^-1-treated *Mettl9^Deg^* NSCs (Supplementary Fig. 5f).

The low levels of residual METTL9 observed in the dTAG^V^-1-treated *Mettl9^Deg^* mESC by WB (see Fig. 3d), as well as the transient presence of the protein from the time of its translation to its degradation, might be sufficient to sustain most of its biological functions in neural cells, resulting in the milder phenotypical and transcriptomic changes of dTAG^V^-1-treated *Mettl9^Deg^* lines compared to the *Mettl9^KO^*. Since the best characterised molecular activity of METTL9 is the catalysis of 1MH modification on target proteins containing the H[ANGST]H motif, we measured bulk 1MH levels in the dTAG^V^-1 treated cells (DIV6) and compared them with the paired control (DMSO) as well as with the *Mettl9^KO^* cells. Interestingly, while 1MH was efficiently reduced in the KO cells, we detected no difference in the 1MH levels between the dTAG^V^-1-treated and DMSO (Ctrl) cells (Fig. 3i) and also the METTL9-independent 3MH levels, used as a control, were not affected (Supplementary Fig. 5g). Overall, these data suggest that the low levels of residual METTL9 protein in the dTAG^V^-1-treated *Mettl9^Deg^*line might be sufficient to sustain neural commitment of mESCs either via METTL9-dependent catalytic functions and/or through other non-catalytic activities.

### METTL9 supports neural commitment of mESCs largely independently of its catalytic activity

To investigate the contribution of METTL9 catalytic activity to its overall biological function, we abolished it by using CRISPR/Cas9. To this end, we targeted the endogenous *Mettl9* locus (Exon 3) in mESCs and edited 4 nucleotides to generate substitutions in 2 highly conserved amino acids within the SAM binding domain of the METTL9 protein (D151K and G153R) (Fig. 4a) which are known to abrogate METTL9 catalytic activity^14^. We selected homozygous *Mettl9^CatD^* (“CatD”, standing for Catalytically Dead) and clonal *Mettl9^WT^* mESC lines (Supplementary Fig. 6a) and investigated their ability to sustain neural commitment and differentiation.

**Fig. 4:**
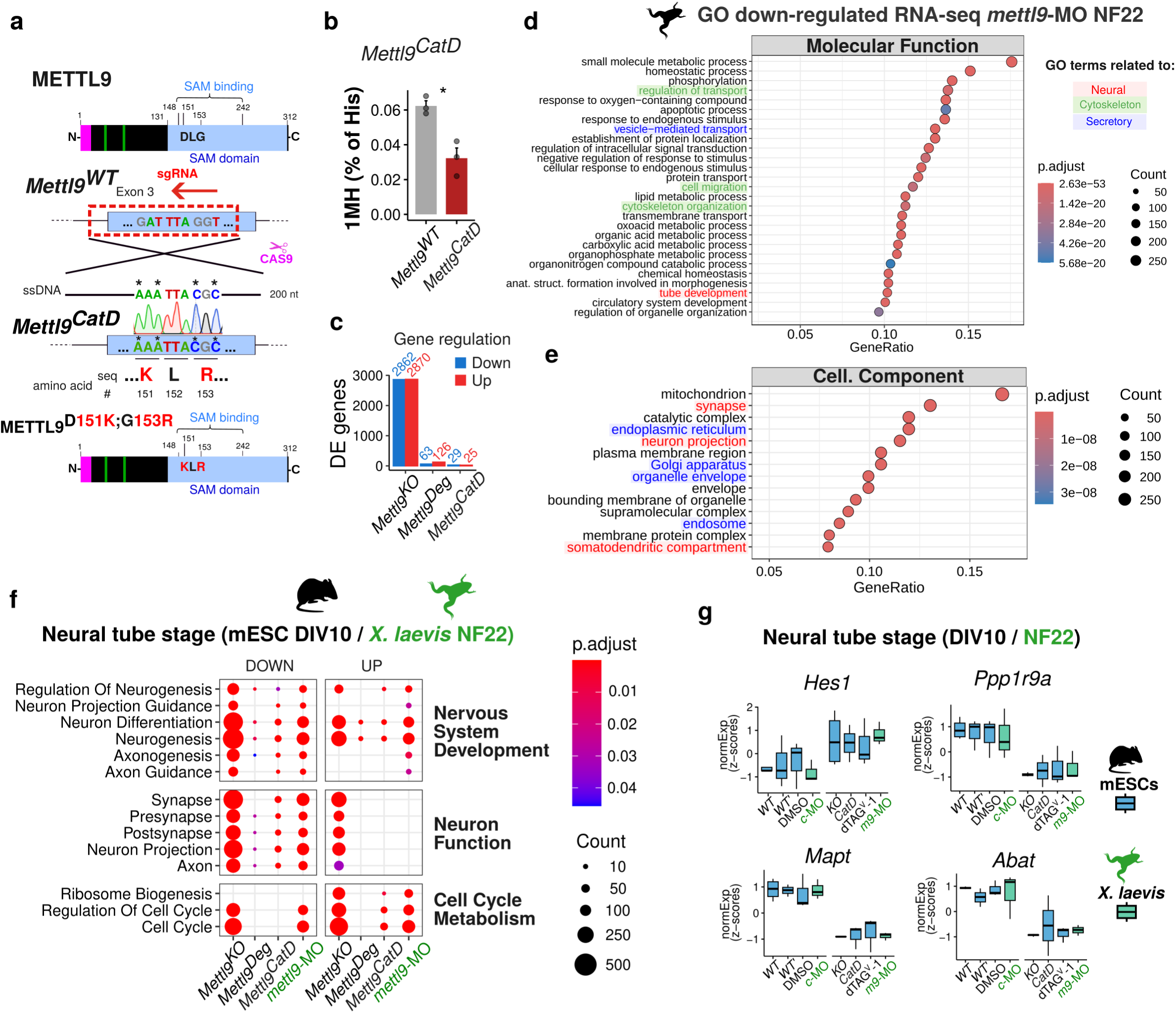
METTL9 catalytic activity only mildly affects neural induction of mESCs. **a** METTL9 protein showing two key catalytic residues (D151, G153) of the SAM binding domain. Below, CRISPR/Cas9 targeting strategy to generate *Mettl9^CatD^* mESCs harboring a mutated METTL9 protein with D151K and G153R. **b** Relative bulk 1MH levels (% of histidines) in *Mettl9^CatD^* and *Mettl9^WT^*NSCs (DIV6) quantified by mass spectrometry (Wilcoxon test). **c** Number of differentially expressed genes found in the RNA-seq (DIV5) in *Mettl9^KO^*, *Mettl9^Deg^* and *Mettl9^CatD^* lines. **d** Top down-regulated Molecular function GO terms in *mettl9*-MO-NF22 *X. laevis* embryos. **e** Top down-regulated Cellular Component GO terms in *mettl9*-MO-NF22 embryos. **f** Neuronal and Cell cycle/Metabolism-related GO terms commonly misregulated among *Mettl9* depleted NPCs (DIV10) and NF22 embryos. ***g*** Example of key neural marker genes consistently misregulated among the different datasets, included in the GO terms in panel f. *WT’* is the clonal WT control for *Mettl9^CatD^*. *c-*MO and *m9*-MO are *ctrl*-MO and *mettl9*-MO respectively.

Surprisingly, *Mettl9^CatD^* NSCs did not display any major macroscopical defect, as shown by comparable NESTIN expression of both *Mettl9^WT^* and *Mettl9^CatD^* lines (Supplementary Fig. 6b), similarly to *Mettl9^Deg^* cells (Supplementary Fig. 5f). To understand whether *Mettl9^CatD^* NSCs exhibited any alteration at the molecular level, we characterised *Mettl9^CatD^*transcriptome by RNA-seq: at DIV5, we found only 54 genes mis-regulated compared with the clonal *Mettl9^WT^* line (29 down-regulated and 25 up-regulated) (Supplementary Fig. 6c; Supplementary Data 1). To exclude that the absence of an appreciable neural phenotype and the mild transcriptomic alterations were due to METTL9 residual catalytic activity in *Mettl9^CatD^*NSCs, we analysed bulk 1MH levels in *Mettl9^CatD^* at DIV6. *Mettl9^CatD^*NSCs displayed a significant reduction in 1MH levels (Fig. 4b), comparable to the 1MH decrease observed in *Mettl9^KO^* NSCs (Fig. 3i), despite having normal levels of the METTL9-independent 3MH modification (Supplementary Fig. 6d). This reduction is in agreement with previous data on METTL9-CatD in HEK293T cells^14^ and indicates that *Mettl9^CatD^*mESCs express a *bona fide* catalytically dead METTL9 protein.

The mis-regulated genes found in the *Mettl9^CatD^* lines represent a much smaller proportion of those previously found in the *Mettl9^KO^* and also smaller than the dTAG^V^-1-treated *Mettl9^Deg^* lines at the same developmental stage (DIV5) (Fig. 4c). Overall, considering also 1MH levels in the 3 cell lines, these data indicate a lack of correlation between the molecular phenotype and the METTL9 catalytic activity (as evaluated by the reduction in 1MH levels). Conversely, METTL9 levels themselves (regardless of the presence of a WT or CatD protein) correlate with the severity of the phenotype; indeed, *Mettl9^KO^* lines which do not produce any METTL9 protein display a massively altered transcriptome; dTAG^V^-1-treated *Mettl9^Deg^* have residual METTL9 protein with normal 1MH levels and relatively mild transcriptomic changes; whereas *Mettl9^CatD^* have a mutant METTL9 protein without catalytic activity, and very mild transcriptomic alterations.

We next analysed the long-term consequences of METTL9 depletion or loss of catalytic activity by performing RNA-seq at DIV10 on *Mettl9^Deg^* (DMSO-, for Ctrl, and dTAG^V^-1-treated) and clonal *Mettl9^WT^* and *Mettl9^CatD^* NPCs (Supplementary Fig. 6e,f; Supplementary Data 1). Consistent with our observations at DIV5, both dTAG^V^-1-treated *Mettl9^Deg^* and *Mettl9^CatD^* cell lines displayed much milder transcriptomic alterations in terms of number and strength of mis-regulated genes compared to *Mettl9^KO^*NPCs.

Interestingly, when comparing the number of mis-regulated genes between *Mettl9^CatD^* and *Mettl9^WT^* NPCs, we found a higher number of differentially expressed genes (DEG) at DIV10 compared to DIV5 (271 versus 54 respectively; Supplementary Fig. 6f,c). This indicates that the loss of METTL9 catalytic activity becomes more detrimental at later stages of neurogenesis, compared to the early phases of neural commitment. Conversely, dTAG^V^-1-treated *Mettl9^Deg^* NPCs (when compared to their paired DMSO-treated cells) displayed milder transcriptomic alterations at DIV10 (63; Supplementary Fig. 6e) than at DIV5 (189; Supplementary Fig. 5b), possibly due to compensatory mechanisms and/or differing molecular roles and interactions of the METTL9 protein during neural differentiation.

Given the striking neuralization defect of *Mettl9^KO^*mESCs at later stages of *in vitro* differentiation, we sought to complement and validate our observations in *Xenopus* embryo. Therefore, we profiled gene expression in whole *mettl9*-MO *Xenopus* tailbuds at stage NF22 (i.e. after neural tube formation), a developmental timepoint roughly comparable to mouse NPCs at DIV10. Interestingly, we found 1617 down-regulated genes and 1454 up-regulated genes in the *mettl9*-MO embryos compared to *ctrl*-MO (Supplementary Fig. 7a). GO analysis revealed many Molecular Functions being down-regulated such as “tube development” as well as “vesicle-mediated transport”, “protein transport”, “cell migration” and “cytoskeleton organization” (Fig. 4d). Among the Cellular Component terms “synapses”, “neuron projection”, and “somatodendritic compartment” as well as “ER”, “plasma membrane region”, “Golgi” and “organelle envelope” were significantly down-regulated (Fig. 4e; Supplementary Data 2). Similarly to *Mettl9^KO^*NPCs, *mettl9*-MO embryos also showed up-regulation of many metabolic genes and interestingly, “central nervous system development” term (Supplementary Fig. 7b).

In conclusion, upon *mettl9* loss, key components linked to neural development are negatively affected at NF22 at a whole-embryo level.

Despite the evolutionary distance, we found many ontologies related to neurogenesis (Fig. 4f) and key neuronal genes consistently mis-regulated both across mouse RNA-seq datasets (*Mettl9^KO^*, dTAG^V^-1-treated *Mettl9^Deg^* and *Mettl9^CatD^*) at DIV10, and in NF22 *mettl9*-MO *Xenopus* embryos (Fig. 4g and Supplementary Fig. 7c). These data strongly support the notion that METTL9 is required throughout vertebrate neural development. However, the milder effects due to either the decreased bulk protein levels in *Mettl9^Deg^* or the specific ablation of the enzymatic activity in *Mettl9^CatD^* suggest that: (i) low levels of METTL9 are mostly sufficient to support neurogenesis and that (ii) this role occurs mainly through catalytic-independent functions.

### METTL9 modulates the secretory pathway in mouse NSCs

We next set out to investigate the molecular functions of METTL9, in NSCs. To this end, we first employed the *Mettl9^Deg^* cell line. Although this system does not achieve a complete protein depletion, it enables rapid and inducible degradation of endogenous METTL9. With this approach we sought to characterise the immediate and direct consequences of METTL9 depletion, which might point at the specific biological roles exerted by METTL9 in NSCs. We characterised the proteome of NSCs by mass spectrometry after only 48 hrs of dTAG^V^-1 treatment, at the onset of neural commitment (from DIV3 until DIV5) (Fig. 5a) and found 2 proteins significantly down-regulated and 11 up-regulated (as compared to DMSO-treated samples; adjusted p value < 0.05) (Fig. 5b and Supplementary Fig. 8a; Supplementary Data 3). These included the ER-to-Golgi and intra-Golgi trafficking protein USO1^40–42^, the ER– and lipid-droplet-associated NSDHL enzyme involved in cholesterol biosynthesis^43^, the plasma membrane-associated FXYD6 (sodium/potassium ATPase regulator)^44^ and carboxypeptidase D, CPD^45^, the ER-associated CYP51a1 involved in cholesterol biosynthesis^46,47^ and the ER associated RDH11 retinol dehydrogenase^48^, all associated with the endomembrane compartment of the secretory pathway. GO analysis on the mis-regulated proteome (encompassing a larger set of 115 up-regulated and 63 down-regulated proteins with a less stringent q value threshold of 0.2) revealed up-regulation of terms related to “ER membrane”, “ER lumen” and “ER protein-containing complexes” (among which chaperones) and “ERGIC” (ER Golgi intermediate compartment) (Fig. 5c) as well as down-regulation of ribosomal-related genes (Supplementary Fig. 8b). We also found up-regulation of terms related to the “Golgi stack” and “Golgi membrane” and “Golgi associated vesicles”, as well as “COPI-coated vesicles” and “exocytic”, “transport” and “secretory vesicles” terms (including synaptic ones) (Fig. 5c). In summary, the proteomic analysis of NSCs after acute METTL9 loss reveals the up-regulation of the endomembrane/secretory pathway as an early molecular consequence.

**Fig. 5:**
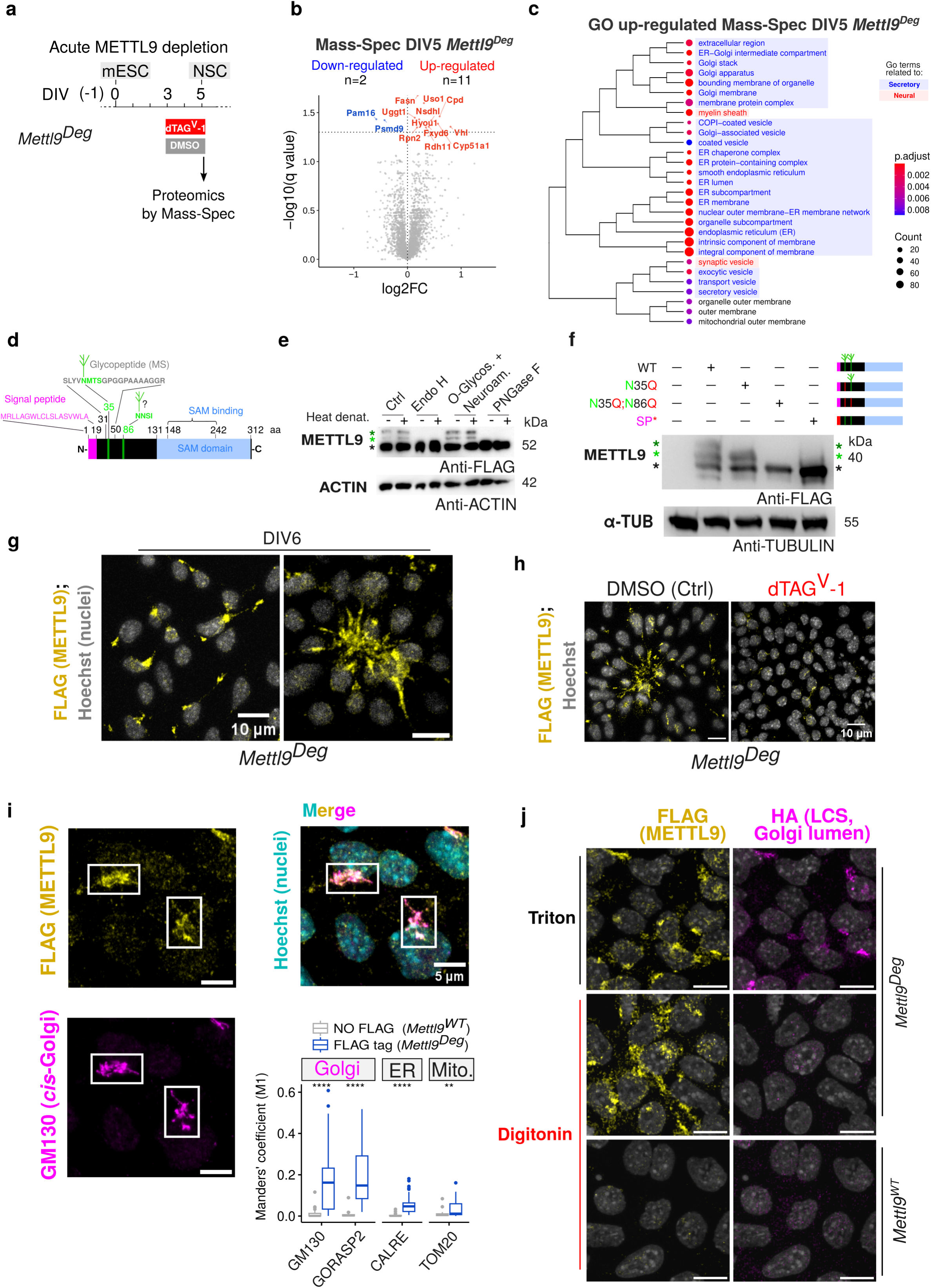
METTL9 is associated with the secretory pathway, and it co-localises with the peripheral Golgi. **a** Scheme depicting the experimental strategy for acute METTL9 depletion in *Mettl9^Deg^*mESCs. **b** Volcano plot of the mis-regulated proteins (coloured dots and labels) in dTAG^V^-1-treated *Mettl9^Deg^* NSCs over Ctrl (DMSO-treated), identified by mass spectrometry (q value < 0.05). **c** GO terms of the up-regulated proteins (with a q value <0.2). The clustering represents all significantly enriched GO terms with an over-representation analysis q value <0.05. **d** METTL9 protein showing signal peptide (magenta), the glycopeptide position^53^ (grey), within sequon (green) and the predicted sequon at position N86 (aa is amino acid). **e** WB showing METTL9-DEG (anti-FLAG antibody) in N-(Endo H and PNGase) or O-glycosydase-treated mESCs extracts. Black, green and dark green asterisks (*) refer to the lowest, intermediate and top METTL9 bands, respectively. **f** WB showing WT METTL9-FLAG or mutated METTL9: METTL9-N35Q-FLAG (N35Q), METTL9-N35Q;N86Q-FLAG (N35Q;N86Q), and SP*-METTL9-FLAG (SP*) (SP* is a mutated Signal peptide), expressed in WT ESCs. In green, N-glycosylated residues; in red, mutated amino acid residues; in magenta the SP. Anti-FLAG and anti-alpha-TUBULIN antibodies were used respectively for METTL9 and loading control. Green and black asterisks near the blot as in (**f**). **g** Representative IF images of *Mettl9^Deg^*NSCs stained with anti-FLAG antibody for METTL9 and Hoechst (nuclei). On the right, a neural rosette is shown. Scale bar is 10 μm. **h** Representative IF images of DMSO-(Ctrl) or dTAG^V^-1-treated *Mettl9^Deg^* NSCs stained with anti-FLAG antibody showing METTL9. Scale bar is 10 μm. **i** Representative IF images of *Mettl9^Deg^* NSCs stained with anti-FLAG antibody (yellow) for METTL9 and GM130 (magenta) for *cis*-Golgi. Scale bar is 5 μm. On the right, co-localisation analysis between anti-FLAG and the following markers: (i) anti-GM130, (ii) anti-GORASP2 (i and ii for Golgi), (iii) anti-CALRE (for ER) or (iv) anti-TOM20 (for mitochondria). The boxplot shows the Manders’ coefficient (M1) values indicating the co-localization between METTL9-FLAG signal and the different subcellular compartment markers. (0<M1<1; total number of slices analysed in *Mettl9^WT^*, NO FLAG, and *Mettl9^Deg^*, FLAG tag: 32, 31 respectively for CALRE; 35 and 40, for GM130; 35 and 31 for GORASP2; 30 and 30 for TOM20. Wilcoxon test). **j** Representative IF images of *Mettl9^Deg^*/LCS-HA or WT E14 NSCs (DIV5), after Triton or digitonin permeabilization. Cells were co-stained with an anti-FLAG antibody (METTL9, yellow), anti-HA (LCS, *trans*-Golgi lumen, magenta) and Hoechst (nuclei, gray). Scale bar is 10 μm.

To investigate whether similar biological processes were still affected after a complete and long-term METTL9 depletion, we characterised the proteome of *Mettl9^KO^* NSCs by mass spectrometry. Interestingly, we found 395 proteins significantly down-regulated and 329 up-regulated over the Ctrl *Mettl9^WT^* cells (q value < 0.01) (Supplementary Fig. 8c; Supplementary Data 3), highlighting a much stronger proteome mis-regulation compared to that occurring in the dTAG^V^-1 *Mettl9^Deg^*line (34 proteins). This is consistent with a complete and prolonged absence of METTL9 during NSC differentiation. Interestingly, GO analysis of the mis-regulated proteins in the KO revealed down-regulation of terms related to “plasma membrane”, “ER”, “ER-Golgi intermediate compartment”, as well as “nervous system development” among others (Supplementary Fig. 8d). These GO terms are consistent with those found at the transcriptomic level in the same *Mettl9^KO^* NSCs (see Fig. 2j), and also with an impaired neuralisation. Among the up-regulated GO were found terms related to “mitochondrion”, “cellular response to leukaemia inhibitory factor”, “endosome membrane”. We then evaluated whether the proteins perturbed in the dTAG^V^-1 *Mettl9^Deg^* were similarly altered in the *Mettl9^KO^*: interestingly we observed that the up-regulated proteins in the dTAG^V^-1 were significantly down-regulated in the *Mettl9^KO^*, and the down-regulated proteins in the dTAG^V^-1 were significantly up-regulated in the *Mettl9^KO^*(Supplementary Fig. 8e). These data indicated that the secretory pathway-related proteins are affected in both systems, although they are regulated in opposite directions and to a different extent (number of proteins and strength of the regulation). Therefore, the alteration of the secretory pathway could be an early cellular event directly linked to METTL9 acute loss, which is exacerbated in the long-term (*Mettl9^KO^*), where complex indirect and/or compensatory cellular mechanisms might also be put in place by NSCs.

### METTL9 localises to the Golgi in mouse NSCs

METTL9 contains a predicted N-terminal signal peptide (SP)^49,50^ corresponding to the first 18 amino acids (Supplementary Fig. 8f). Such peptide sequences target proteins to the secretory pathway via the ER^51^. Western blotting of both the endogenous wild-type METTL9 (Fig. 2c) and METTL9-DEG (Fig. 3c) proteins highlighted the existence of multiple METTL9 bands, suggesting the presence of post-translational modifications in both mNSCs and mESCs. METTL9 contains a canonical NMTS glycosylation sequon (amino acid # 35-38)^52^ (Fig. 5d, green) with the Asn (N35) as a possible glycosylation site^53^*. In silico* prediction analysis^54^, revealed N86 as an additional putative N-glycosylation site (Fig. 5d), although with lower probability than N35. We thus sought to determine whether the shifted METTL9 bands observed by SDS-PAGE are due to protein glycosylation in mESCs. We treated protein extracts with different glycosidases^55^ *in vitro* and found that the 2 highest METTL9 bands completely disappeared upon N-glycosidase (Endo H or PNGase F) treatment, but not in the control or in O-glycosidase-treated samples (Fig. 5e). Given the sensitivity to Endo H^55^, these data suggest that the two highest METTL9 bands corresponded to two high-mannose (or non-complex hybrid) N-glycans.

To directly prove this, we generated two mutagenised constructs encoding for a METTL9-FLAG coding sequence harbouring either one amino acid substitution (N35Q) or two (N35; N86Q), which prevent N-glycosylation (Fig. 5f and Methods), and transfected them in mESCs. Despite the presence of 3 bands in the control, in the N35Q mutant the highest METTL9 band was lost, whereas in the double mutant (N35Q;N86Q) only the lower METTL9 band was observed (Fig. 5f). This confirms that both N35 and N86 are N-glycosylated in mESCs. In addition, we generated a fourth construct to alter the tripartite regions^56,57^ of METTL9 signal peptide (SP*-METTL9-FLAG: R2A;W7A;C9A;S11A; see Methods). Although no N-glycosylation site was edited in this construct, it showed only the lowest, unmodified METTL9 band by WB, similarly to the double mutant (N35Q;N86Q) (Fig. 5f and Supplementary Fig. 8g). This result suggests that disruption of the SP prevents the N-glycosylation of both Asn residues (N35 and N86), most likely by precluding METTL9 translocation into the ER.

These data were further corroborated through the inhibition of the first biosynthetic step of N-linked glycosylation in the ER by supplying tunicamycin^58–60^ to mESCs: protein extracts were analysed by WB, which revealed the complete depletion of the two highest METTL9 bands in tunicamycin-treated samples (Supplementary Fig. 8h).

Overall, our experiments suggest that METTL9 N-terminal SP directs the protein to the ER lumen, where it acquires high mannose N-glycans on two distinct Asn residues (N35;N86). Indeed, this is also supported by available proteomic data^61^, where METTL9 was found to co-fractionate mainly with the ER and other compartments of the secretory pathway in 5 different human cell lines (Supplementary Fig. 8i). Furthermore, since METTL9 lacks typical ER retention signals, such as KDEL^62^, it likely exits the ER and proceeds further along the secretory pathway, through the Golgi.

To further validate METTL9 association with the secretory pathway, we determined its endogenous sub-cellular localisation. Due to the lack of suitable antibodies, we took advantage of the FLAG tag of the *Mettl9^Deg^*cell line and performed immunofluorescence staining of NSCs: this revealed that almost all cells expressed METTL9 (Fig. 5g, left panel), indicating that METTL9 is homogenously expressed in NSCs.

Moreover, METTL9 displayed a distinct, asymmetrical cytosolic distribution (Fig. 5g), particularly evident in the apical part of neural rosettes (Fig. 5g, right panel); this signal was greatly reduced upon dTAG^V^-1 treatment (Fig. 5h), highlighting the specificity of the FLAG antibody in recognising the METTL9 protein. Since METTL9 distribution was reminiscent of the Golgi apparatus, which is asymmetrically localised in neural cells^63–66^ and in particular within neural rosettes^67,68^, we assessed whether METTL9 co-localised with it. Indeed, co-staining of NSCs with the *cis-*Golgi markers GM130^69^ and GORASP2^70^ confirmed an extensive overlap between METTL9 and the Golgi (Fig. 5i and Supplementary Fig. 8j) in DIV6 cells, whereas a very low proportion of METTL9 signal was found to co-localise with other cytosolic organelle markers such as CALRE for the ER^71^ or TOM20 for mitochondria^72^ (Fig. 5i and Supplementary Fig. 8j). Consistently, METTL9 subcellular distribution drastically changed in DIV6 NSCs after an hour treatment with golgicide (Supplementary Fig. 8k), a drug which specifically induces Golgi fragmentation^73^. Overall, these data indicate that METTL9 is a Golgi-associated protein in mouse NSCs, whose subcellular localisation depends on the integrity of the Golgi.

Eventually, we assessed whether METTL9 localisation at the Golgi occurred on its peripheral (cytosolic) or luminal side (or both). To test this, we performed immunofluorescence on NSCs after selective plasma membranes permeabilization with digitonin, which leaves cholesterol-poor membranes, such as that of Golgi, intact^74,75^ and thus not accessible to antibodies. A tagged-version of the *trans*-Golgi Lactosylceramide synthase (LCS-HA) was used as a Golgi luminal control (see Methods). As expected, the LCS-HA signal was detected only in the Triton-NSC samples (control), but not in the digitonin one (Fig. 5j); on the contrary, the FLAG signal of the endogenously tagged METTL9 was detected in the digitonin-permeabilised sample and comparable to the Triton-permeabilised, indicating that METTL9 is positioned on the cytosolic face of the Golgi.

Overall, this data showed a tight association of METTL9 with the secretory pathway since i) it is N-glycosylated in this compartment, ii) it localises to the peripheral (cytosolic) Golgi side and iii) it modulates the secretory pathway-related proteome. This suggests that METTL9 could exert novel molecular functions linked to the homeostasis of this cellular compartment.

### METTL9 interacts with key secretory pathway and transport regulators, independently of its catalytic activity

To investigate the molecular pathways regulated by METTL9, we characterised the METTL9 interactome in NSCs (DIV4), by performing immunoprecipitation (Supplementary Fig. 9a) coupled to mass spectrometry (IP-MS; Supplementary Fig. 9b). This identified 71 proteins enriched in the Anti-FLAG-METTL9 IP (Fig. 6a; Supplementary Data 3). Among the top interactors were the microtubule destabiliser Stathmin1, (STMN1), which is highly expressed in the nervous system also during development^76–78^ and the E3-ufmylation adapter DDRGK-domain containing protein 1, DDRGK1, anchored to the cytosolic side of the ER^79^ and involved in reticulophagy^80^. Gene Ontology (GO) analysis (Fig. 6b) revealed that METTL9 interactome is enriched in Golgi membrane, ER protein-containing complexes, synaptic vesicles, transport vesicles and lysosomal and endosomal membrane factors. Importantly, beside other know METTL9 interactors like CANX and FAF2, we found many RAB proteins, including the pre-Golgi and cis-Golgi RAB1A and RAB2A (both involved in pre-Golgi trafficking and whose knock-down or over-expression cause Golgi fragmentation^81–84^), and the late-endosome and lysosomal GTPase RAB7^85,86^. We validated METTL9-STMN1 and METTL9-RAB2a physical interactions after co-expressing METTL9-FLAG and either STMN1-HA or HA-RAB2a in mESCs. METTL9 was co-immunoprecipitated by STMN1-HA or HA-RAB2a, as shown by WB (Fig. 6c, d).

**Fig. 6:**
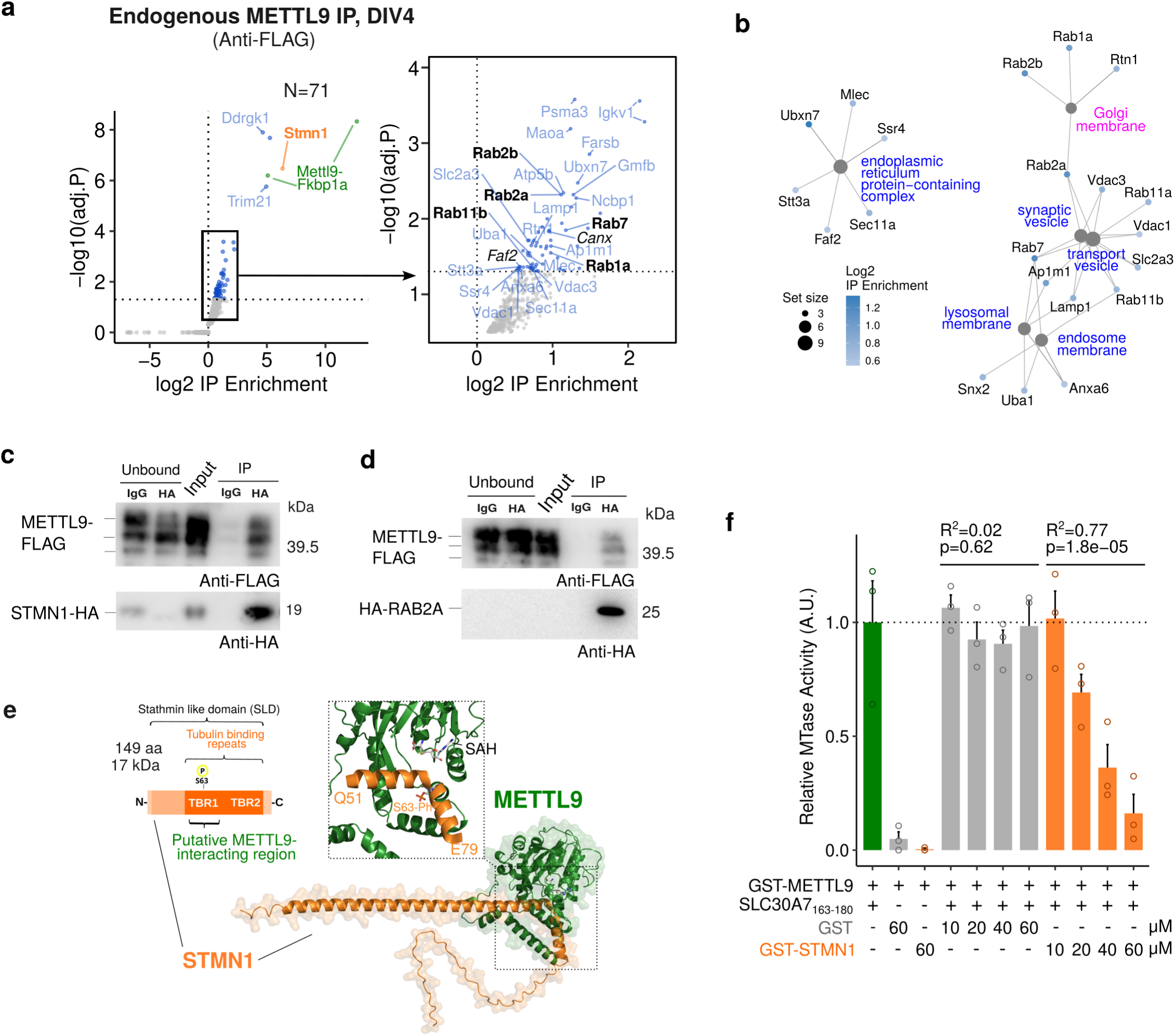
METTL9 interacts with secretory pathway-related proteins in mNSCs. **a** Volcano plot showing the proteins enriched upon METTL-IP (anti-FLAG) over the control-IP (IgG) in *Mettl9^Deg^* NSCs (q value < 0.01); on the right, zoom-in on the enriched interactors, among which the known METTL9-interactors *Faf2* and *Canx* (in black, italics). **b** Network showing the genes belonging to the top GO terms enriched in METTL9 interactors relative to the secretory pathway. **c, d** WB showing the immunoprecipitation of STMN1-HA (IP) (c) or HA-RAB2a (d) with anti-HA beads (HA) or IgG (ctrl), after co-expression of STMN1-HA or HA-RAB2a and METTL9-FLAG in WT mESCs. WB signal is detected by anti-FLAG and anti-HA. **e** AlphaFold modelling prediction of STMN1-METTL9 protein complexes (STMN1, in orange; METTL9 in green). **f** Bar graph showing *in vitro* METTL9 methyltransferase activity (MTase) of recombinant GST-METTL9-FLAG (GST-METTL9) protein with the SLC30A7_163-180_ peptide or with GST-STMN1-HA (GST-STMN1) and GST (first, second and third bar, respectively); GST-METTL9 activity was also measured with increasing concentrations (μM) of GST or GST-STMN1. The correlation between MTase activity and concentration of GST or GST-STMN1 is expressed by the R^2^ and p values (linear regression).

To further characterise the METTL9-STMN1 interaction, we predicted the structure of this complex *in silico* by using AlphaFold^87^ (Fig. 6e). Interestingly, the METTL9 amino acid residues mainly involved in the binding coincide with its catalytic pocket (Fig. 6e). It is noteworthy that STMN1 amino acid sequence, and in particular the residues predicted to participate in METTL9 binding contain no *bona fide* 1MH motif and thus are unlikely to be methylated. Overall, these data suggest that METTL9 might engage STMN1 in a catalytic-independent interaction.

We tested these predictions by measuring METTL9 methyltransferase activity (MTase) in a biochemical assay (Supplementary Fig. 9c,d), in the presence of *S*-adenosylmethionine (SAM) and the SLC30A5_163-180_ synthetic peptide, which is a known METTL9 target^14^. When using purified recombinant GST-STMN1 (GST-STMN1-HA) as a substrate instead of SLC30A5_163-180_, GST-METTL9 MTase activity was almost undetectable, and comparable to the negative GST control (Fig. 6f). This strongly supports the notion that STMN1 is not a METTL9 substrate *in vitro*. We next assessed METTL9 MTase activity on the canonical SLC30A5_163-180_ target peptide, in the presence of increasing concentrations of STMN1. Remarkably, we observed that METTL9 activity was impaired by STMN1 (but not by GST) in a dose dependent manner (Fig. 6f). These data indicate that STMN1 is able to outcompete a known METTL9 substrate *in vitro* and are in agreement with our structural models showing STMN1 in the catalytic pocket of METTL9.

AlphaFold predictions also showed that METTL9 embraces a few amino acids of the STMN1-Tubulin binding repeat I, including Ser63 (Fig. 6e), whose phosphorylation regulates tubulin binding^88–90^. Therefore, METTL9 binding may modulate STMN1 function, similarly to the way in which STMN1 regulates METTL9 catalytic activity. A comparable mechanistic model might be extended to other members of the METTL9 interactome, like RAB2A and RAB7A (Supplementary Fig. 9e,f), which share with STMN1 i) a predicted binding to the METTL9 catalytic pocket, ii) the lack of H[ANGST]H motifs and iii) the presence of regulatory domains (i.e. switch II GTPase domain) contacting METTL9. Thus, similarly to STMN1, also the activity of these RAB proteins could be potentially regulated upon METTL9 binding.

Interestingly, the substitutions D151K and G153R within METTL9-CatD are predicted to have a minimal impact on the overall METTL9 protein structure (Supplementary Fig. 9g,h). The mutated residues are in close proximity to the most buried and solvent-inaccessible end of the catalytic site, where the SAM molecule is accommodated. Notably, they are positioned far from the bulk of the predicted interaction surface between METTL9 and STMN1 or RAB2A (Supplementary Fig. 9i), suggesting that METTL9 could preserve these protein-protein interactions regardless of its catalytic activity. Consistent with these models, METTL9-CatD-FLAG could be immunoprecipitated by STMN1-HA or HA-RAB2a, after an Anti-HA-IP in mESCs, as shown by WB (Supplementary Fig. 9j,k, respectively).

Overall, the fact that the protein-protein interactions between the catalytically inactive METTL9 and STMN1 or RAB2a are likely preserved in *Mettl9^CatD^* mESCs might explain why they show a mild molecular phenotype and highlights the importance of these physical interactions in NSCs.

### The catalytic-dependent function of METTL9 has a secondary but convergent role in neural development and secretory system function

METTL9 catalytic activity exerts a secondary role compared to the main non-catalytic functions in NSCs, as shown by the less severe neural differentiation defects of *Mettl9^CatD^* compared to *Mettl9^KO^*. In agreement with this, there is a striking difference between *Mettl9^KO^*and *Mettl9^CatD^* or *Mettl9^Deg^* in the number and strength of mis-regulated genes. However, the three cell lines also share a high overall level of consistency among their transcriptomic alterations. Indeed, cumulative plots of the Fold Changes (Supplementary Fig. 10a) showed that many hundreds of the most up-(or down-) regulated genes in the *Mettl9^KO^* were consistently up-(or down-) regulated, respectively, in the other lines, although to a milder extent. In addition, integrative re-analysis of all the three mESC experiments at DIV5 confirmed “neurogenesis”, “neuron/cell projections”, “synapses”, “somatodendritic” compartment and “Golgi” terms among the top gene ontologies coherently down-regulated in all the cellular models (Supplementary Fig. 10b,c). Moreover, we found that many genes encoding for Golgi-resident enzymes or structural Golgi-proteins; transport– and secretory pathway-related proteins (also involved in neural processes) were consistently mis-regulated, albeit to different extents, among all mouse RNA-seq datasets (*Mettl9^KO^*, *Mettl9^Deg^*, *Mettl9^CatD^*) at DIV5 (Supplementary Fig. 10d).

Therefore, notwithstanding the less severe neural phenotype of the *Mettl9^CatD^*, the high degree of similarity of the affected cellular pathways among the different cell lines suggests that 1MH-dependent and independent activities might converge onto the same molecular processes in NSCs. A prerequisite for this model would be that METTL9 substrates of methylation might be directly involved in secretory-related and neuronal pathways. To test this hypothesis, we scanned the sequences of the mouse proteome and identified potential METTL9 substrates (i.e. all H[ANGST]H-containing proteins). Interestingly, GO analysis revealed that secretory, neuronal and transport-related processes accounted for almost 50% of the categories significantly enriched in putative 1MH targets (Supplementary Fig. 10e). Moreover, some of these proteins have already been demonstrated to be methylated by METTL9 in HEK293T cells (Supplementary Fig. 10f)^14^. These included MYO18A, known to exert important roles in Golgi positioning in neural cells^65^ and many zinc transporters (SLC30A1/5/7 and SLC39A7), which are critical for controlling zinc levels within the cell, and some also within organelles such as the Golgi compartment^91,92^. Therefore, while METTL9 has potentially hundreds of targets expressed in any cell type, their 1MH-modification (and hence regulation) might be more critical for neural stem cells, which heavily rely on the secretory pathway to sustain directional trafficking towards the apical part of the cell, where the growth cone emerges^93,94^.

Overall, these data suggest that, while the enzymatic and catalytic-independent functions of METTL9 act through distinct molecular mechanisms (1MH-methylation of protein substrates and protein-protein interactions, respectively) and that their relative contribution greatly varies in NSCs, both functions might impinge on the same cellular machinery of the secretory system, sustaining proper neural development.

### METTL9 depletion affects cellular trafficking kinetics and disrupts Golgi integrity in mNSCs

We showed that the expression levels of hundreds of early secretory pathway proteins was altered in *Mettl9^KO^* NSCs (Supplementary Fig. 8c,d). A similar (although to a much smaller extent) effect was observed upon METTL9 acute depletion (Fig. 5b,c); this suggested that the homeostasis of the secretory pathway could be directly controlled by METTL9. Moreover, many METTL9 interactors control key cellular processes such as macromolecular motility and cargo engagement; for instance, STMN1 controls microtubules growth^76,77^ and cellular trafficking along the secretory system also relies on this cytoskeleton component^95^. RAB2a is a key regulator of ER to Golgi trafficking and its GTPase domain is potentially modulated by METTL9 binding (Supplementary Fig. 9e,f).

Thus, we investigated the ER to Golgi trafficking kinetics of a Golgi resident enzyme, the α-mannosidase II (ManII), by using the retention using selective hooks (RUSH) method^96^, in *Mettl9^KO^*NSCs (Fig. 7a). This state-of-the-art system consists of two components, a hook and a reporter: at the steady state, the hook, which is an ER-localised Streptavidin protein (Str-KDEL), anchors the cargo, ManII, fused to the streptavidin-binding peptide-EGFP (ManII-SBP-EGFP) to the ER (donor compartment), via the strong Str-SBP interaction. Upon biotin supply to live cells (Time 0, T_0_), the ManII-SBP-EGFP is displaced from the ER-Str hook, enabling its synchronous release from the ER to the Golgi (final, acceptor compartment).

**Fig. 7:**
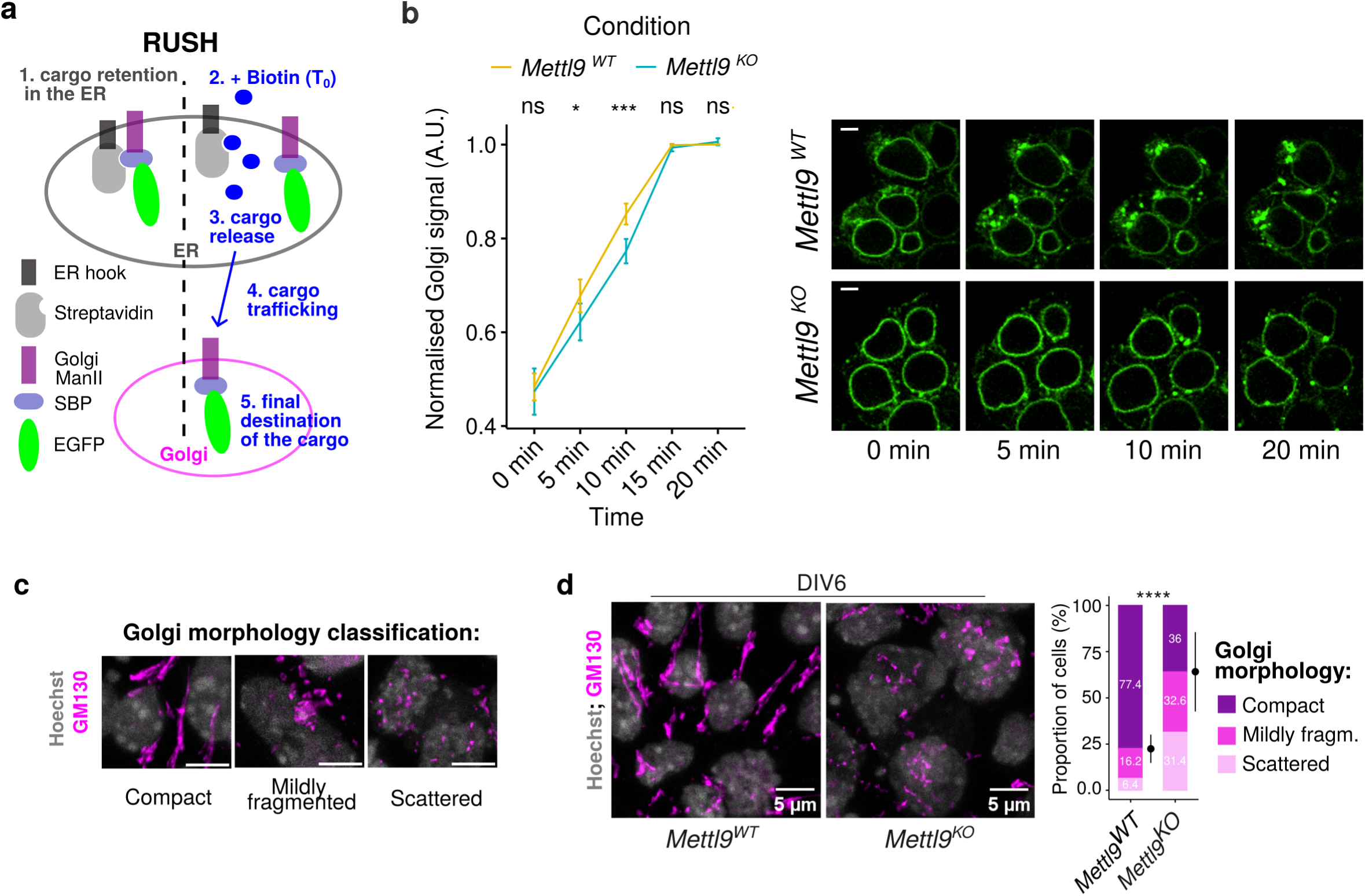
METTL9 depletion impairs cellular trafficking kinetics and Golgi morphology in mNSCs. **a** Schematic showing the RUSH system used to study cellular trafficking in this work. The ER hook here is the Streptavidin protein anchored to the Endoplasmic Reticulum (ER). The reporter includes the streptavidin-binding peptide (SBP) fused to the cargo which is the α-mannosidase II (ManII) enzyme resident in the Golgi and to the fluorescent EGFP protein. Upon Biotin addition to cell media, (T_0_), the reporter is released from the ER hook and its export from the ER starts. Its trafficking is followed until it reaches the Golgi, where ManII is delivered to. **b** Normalised ManII-SBP-EGFP signal in the Golgi of *Mettl9^WT^*and *Mettl9^KO^* NSCs, at different timepoints after Biotin addition (*: P<0.05; ***: P<0.001; T test). On the right, representative close-ups of live microscopy images of MannII-SBP-eGFP signal used for quantification. **c** Classification of Golgi (anti-GM130) morphology in mNSCs into 3 categories, used for cell counting. Scale bar is 5 μm. **d** Representative IF images of *Mettl9^WT^* and *Mettl9^KO^* NSCs stained with anti-GM130 (Golgi) and Hoechst (nuclei). Scale bar is 5 μm. On the right, corresponding quantifications of Golgi morphology (categorised in compact, mildly fragmented or scattered). For *Mettl9^WT^* and *Mettl9^KO^*, 421 and 389 cells were counted, respectively (χ2 test).

We engineered stable *Mettl9^WT^* and *Mettl9^KO^*cell lines expressing both the ER hook and the ManII-SBP-EGFP cargo and performed a time course live imaging experiment at NSCs stage. At T_0_, EGFP signal was mainly localised at the ER (green ring around nuclei) in both *Mettl9^WT^* and *Mettl9^KO^* cells (Fig. 7b); however, 5 and 10 mins after Biotin addition the EGFP signal was significantly more retained in the ER for the *Mettl9^KO^* compared to the *Mettl9^WT^*cells, which had already started to export ManII-SBP-EGFP to the ER exit sites (i.e. more signal as dots and granules compared to the *Mettl9^KO^*). This defect was fully recovered after 20 minutes (endpoint), as most EGFP signal was detected in the Golgi (dots) both in the *Mettl9^WT^* and *Mettl9^KO^* cells (Fig. 7b and Supplementary Fig. 11a,b,c). Overall, this data indicates a slight but significant delay in the kinetics of the cellular trafficking from the ER to the Golgi in *Mettl9^KO^* NSCs; this is consistent with the altered proteomic data in the same cells and could be explained by the disruption of the regulatory functions normally exerted by METTL9 on STMN1 and RAB2 activities.

Besides regulating cellular trafficking, METTL9 co-localises with the Golgi (Fig. 5i) and many of its protein interactors, particularly, STMN1 and RAB2, are essential for maintaining the structural integrity of the Golgi^82,97^. Thus, we investigated whether the absence of METTL9 could have a detrimental effect on the Golgi apparatus morphology. By performing immunofluorescence staining of *Mettl9^KO^* cells with an anti-GM130 antibody, we classified Golgi morphology in three categories^98^: compact (i.e. with a dense and/or elongated shape typical of neural stem cells), mildly fragmented and completely scattered (Fig. 7c), and we enumerated cells according to these 3 categories. Interestingly, we observed that *Mettl9^KO^* NSCs displayed a significantly higher proportion of scattered and mildly fragmented Golgi compared to *Mettl9^WT^*cells at DIV6 (Fig. 7d).

It is noteworthy that both dTAG^V^-1-treated *Mettl9^Deg^*and *Mettl9^CatD^* NSCs also showed a significant, albeit milder Golgi fragmentation pattern (Supplementary Fig. 11d), further supporting the convergence of METTL9 catalytic dependent and independent functions on the maintenance of the secretory system.

Golgi integrity and positioning presides cell polarization, axon elongation and intracellular trafficking, which are pivotal processes in the induction and maturation of a neuronal cell^63,93,94,99^. In light of this, our data suggest that, upon METTL9 depletion, Golgi morphology and cellular trafficking are negatively affected, probably due to the lack of METTL9 regulatory activity on important interactors like STMN1 and RAB2; these cellular defects, in turn, might prevent neural differentiation of mESCs.

### The catalytic independent roles of METTL9 in the secretory pathway and neural development are evolutionary conserved

To enable the molecular dissection of amphibian NSCs and get insight into the conservation of the molecular pathways affected by *mettl9* loss, we took advantage of *Xenopus* animal caps (a.c.) neuralised with *noggin* mRNA. Thus, we assessed the effect of *mettl9* knock-down on early neural induction by analysing the proteome and transcriptome of a.c. at stage 12.5 (Fig. 8a; Supplementary Fig. 12a). Strikingly, proteomic analysis by mass spectrometry revealed the “Golgi” GO term amongst the most down-regulated biological processes (Fig. 8b; Supplementary Fig. 12b; Supplementary Data 3). Furthermore, RNA-seq analysis of the same samples (Supplementary Fig. 12c; Supplementary Data 1) identified amongst the most down-regulated GO terms “projection organization”, as well as “cell motility” and “vesicle-mediated transport” (Fig. 8c; Supplementary Data 2). Moreover, among the down-regulated cell components, we found the “ER”, “cytoskeleton” and “Golgi” (Fig. 8d; Supplementary Data 2). Interestingly, among the most up-regulated Molecular Function terms we found “generation of neurons”, “neurogenesis” and “neuron differentiation” (Supplementary Fig. 12d). These data strongly suggest that *Mettl9* exerts neurodevelopmental functions by modulating similar pathways (i.e. secretory and cytoskeleton) throughout vertebrates. Given the high level of conservation between *Xenopus* and mNSCs (both in terms of neural phenotype and cellular pathways regulated by METTL9, e.g. Golgi), we leveraged *Xenopus* embryos to evaluate the contribution of METTL9 catalytic activity to neural development *in vivo*. To this end, we co-injected a *Xenopus laevis mettl9^CatD^* mRNA (harbouring the same mutations of mouse *Mettl9^CatD^*) (Supplementary Fig. 12e) with *mettl9*-MO in embryos and assessed the potential recovery of the neural phenotype by analysing *elcR* mRNA expression by WISH. Remarkably, *mettl9^CatD^* injected embryos could partially rescue the neural defects ascribed to *mettl9* knock-down. Indeed, a significantly lower proportion of both *mettl9^WT^* and *mettl9^CatD^* embryos showed a perturbed neuralization pattern when compared to the *mettl9*-MO injected embryos alone, rescuing *elcR* expression in the intermediate (i) stria, as well as in the trigeminal placode (tp) (Fig. 8e,f). Importantly, the extent of the recovery in the *mettl9^CatD^* was comparable to that of *mettl9^WT^*, suggesting that *mettl9^WT^* or *mettl9^CatD^* can similarly rescue the neural phenotype in *Xenopus*.

**Fig. 8:**
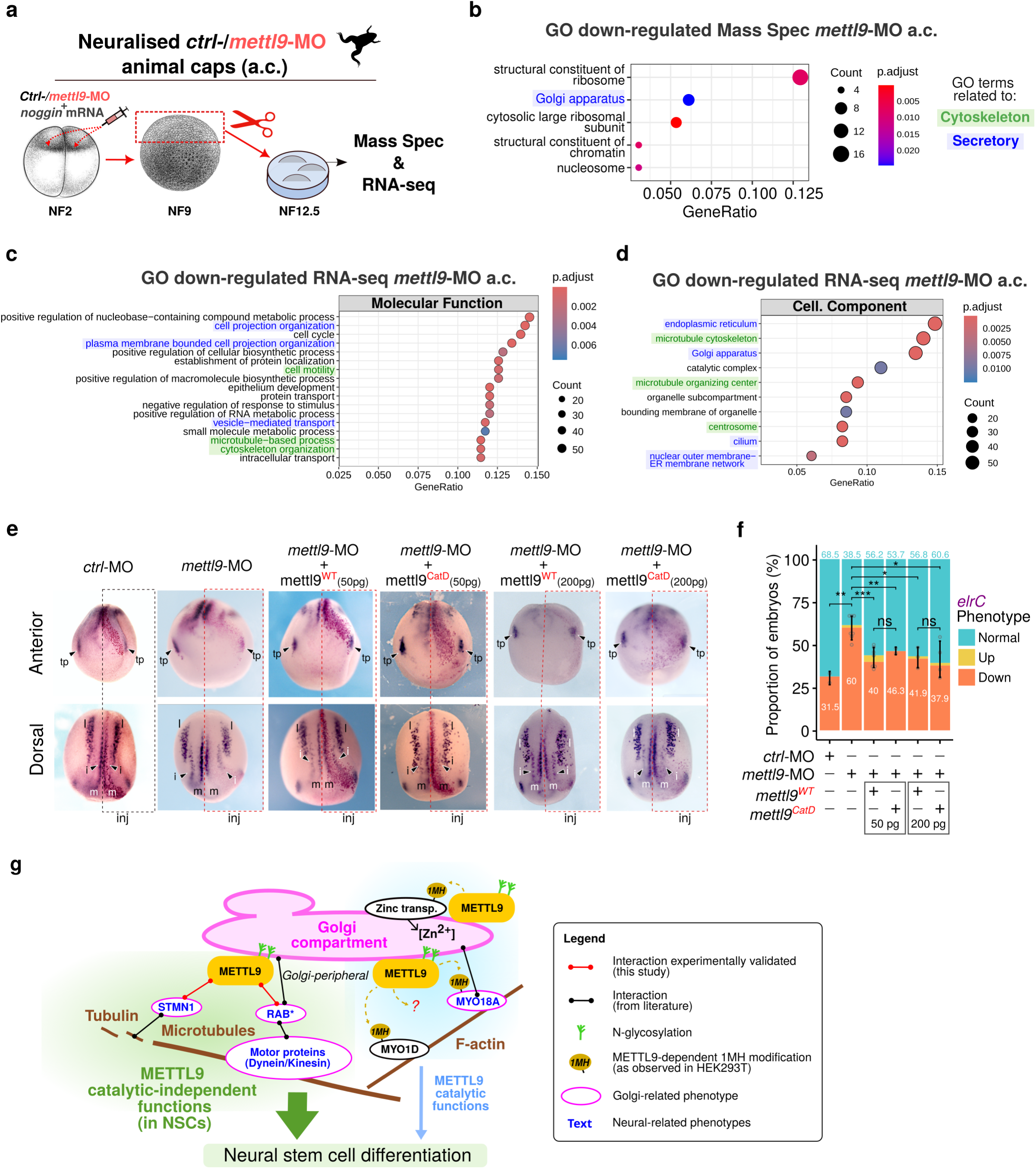
Conserved alteration of the secretory pathway in *mettl9*-MO *X. laevis* neuralised animal caps (a.c.) **a** Schematic depicting the preparation of neuralised animal caps (a.c.) from *mettl9*-MO or *ctrl*-MO embryos in *X. laevis*. Embryo images adapted from Zahn et al.^171^. **b** Most down-regulated GO terms in the *mettl*-MO a.c. proteome. (C-D) Most down-regulated Molecular Function *(C)* and Cellular Component *(D)* GO terms in *mettl9*-MO neuralised a.c. RNA-seq data. GO terms related to Cytoskeleton and Secretory pathways were highlighted in green and blue respectively. **e** Representative anterior and dorsal views images of *X. laevis* embryos injected with either *ctrl*-MO or *mettl9*-MO, or co-injected with *mettl9*-MO and either *mettl9^WT^* or *mettl9^CatD^*mRNAs, showing *elrC* mRNA expression (purple), by whole-mount RNA *in situ* hybridisation. Arrowheads indicate intermediate (i) neuron precursors and trigeminal placodes (tp) affected in the treated side of *mettl9*-MO injected embryos. Inj.: Injected side. (m) medial and (l) lateral stria. **f** Bar graph showing the quantification of embryos: *ctrl*-MO, *mettl9*-MO or *mettl9*-MO co-injected with either *mettl9^WT^* or *mettl9^CatD^*mRNAs, screened for altered *elrC* expression (χ2 test) (percentages of embryos are reported in the graph). **g** Schematic depicting our working model: METTL9 has an evolutionary conserved role in vertebrates in sustaining early neural development, mainly through catalytic independent functions (in green). Among these, we identified one related to the maintenance of the secretory pathway. This function is mediated by protein-protein interactions occurring most likely at the peripheral side of the Golgi (magenta), where METTL9 is localised in mNSCs. We envisage that METTL9 binding to STMN1 and RAB2 regulates their functions related to the cytoskeleton, cargo motility and Golgi structure. Therefore, in METTL9 deficient NSCs, cellular trafficking and Golgi morphology are perturbed, and this is detrimental to the establishment of neural polarity, cell signalling, axon development and ultimately to neural differentiation. METTL9 methyltransferase activity may cooperate with the maintenance of the secretory pathway through histidine methylation (and thus, probable regulation) of Golgi-related substrates, like MYO18A or zinc transporters like SLC30A5/7 and SLC39A7 but might be marginal for neural development.

Overall, these complementation assays confirmed the direct involvement of Mettl9 in neural development *in vivo* and corroborated our previous data about the primary role of Mettl9 catalytic-independent functions in in mNSCs.

## Discussion

In this work, we identified the first developmental role for Mettl9 in the context of neural differentiation. By exploring human and mouse scRNA datasets and using 2 different model systems (namely *X. laevis* embryos and mouse neural stem cells cultures), we showed that Mettl9 is highly expressed during vertebrate neurogenesis. Importantly, we discovered that neural fate specification is consistently impaired by perturbing Mettl9 expression or function with different genetic systems, resulting in aberrant developmental trajectories. Overall, our data demonstrate that Mettl9 requirement for early neurogenesis is a conserved feature of vertebrates.

Our study highlights that this neural function is likely mediated by direct involvement of METTL9 in the secretory pathway: indeed ER-, vesicle– and Golgi-related pathways are mis-regulated upon METTL9 depletion, as shown by proteomic and transcriptomic data, with the latter indicating possible regulatory feedback loops^100–102^. These molecular perturbations are consistent with the altered cellular trafficking kinetics, observed by RUSH, in mNSCs.

Importantly, we found that METTL9 co-localises with the Golgi apparatus in mNSC. We suspect that this might have been overlooked by other workers because of a lack of good anti-METTL9 antibodies, as well as the exclusive use of METTL9 over-expression systems in cancer cell lines.

METTL9 association with the Golgi is *per se* very intriguing, since this organelle underlies multiple aspects of neural fate specification and maturation. Golgi is asymmetrically distributed in neural cells^63–66^ and its position determines neural cell polarity and axonal growth^63^. Indeed, Golgi polarisation underlies asymmetric membrane trafficking, which is key for dendrite and axon growth^93,94,99,103^. In addition, Golgi is also present in the form of outposts within dendrites, where it sustains dendrite growth^99,104^. Therefore, Golgi fragmentation in METTL9 deficient NSCs might have broad consequences on neuronal function, such as disruption of polarised trafficking which in turns could prevent axon growth.

The close association of METTL9 with the secretory pathway is also supported by biochemical data. Indeed, we showed that METTL9, besides harbouring a signal peptide (SP), is modified with high mannose N-glycans in two Asn residues of its amino acid sequence. Thus, METTL9 enters the secretory system through the ER where it is N-glycosylated. Our mass spectrometry data in mNSCs highlighted many METTL9 interactors known to be associated with the peripheral (cytoplasmic) side of Golgi as well as with endosomes and vesicles. This is consistent with METTL9 being detected by IF on the cytosolic side of the Golgi. Interestingly, the METTL9 SP might escape cleavage and could be retained, as a buried helix, within the mature METTL9 protein (Supplementary Fig. 8f). Thus, its SP might act as an anchor to tether METTL9 to the Golgi membrane (similarly to other proteins^105–107^). Alternatively, this localization pattern might be accomplished by specific protein interactors. While our data clearly indicate that METTL9 co-localises with the Golgi apparatus, further work is required to dissect the biological significance of METTL9 N-glycosylation on its catalytic and non-catalytic functions, particularly in the context of neural development.

The focus of METTL9 studies so far has been on its methyltransferase activity^14,18^. The 1MH profiles by Davydova et al.^14^ document particularly high levels of METTL9 activity in the brain. Here, we described that proteins involved in the secretory pathway and Golgi homeostasis are enriched in METTL9 substrates, either experimentally validated^14^ or predicted by the presence of the canonical H[ANGST]H motif. Many zinc transporters are targets of METTL9-dependent methylation (including SLC39A7, SLC30A1/5/7), and the zinc-binding affinity of SLC39A7 is directly regulated by 1MH^14^. These zinc-transporters are embedded into the membrane of the Golgi *cisternae* (except for the plasma membrane-associated SLC30A1), where they regulate zinc transport and concentration^91,108,109^. This function, in turn, is crucial to support the activity of numerous Zn-dependent enzymes of the secretory pathway (e.g. α-mannosidase II and β-4-galactosyltransferase^92,110–112^). The unconventional myosin protein MYO18A is another experimentally validated METTL9 target^14^: it exerts important roles in bridging the Golgi to the actin cytoskeleton via GOLPH3, by shaping Golgi morphology and positioning as well as affecting NSC polarity and apically-targeted membrane trafficking during mouse corticogenesis *in vivo*^65,113–116^. These insights have been highlighted in our study also thanks to the pivotal role of the secretory pathway in highly polarised cells, such as developing neurons.

Notwithstanding the above, we found that METTL9 exerts a significant role independently of its catalytic activity. By comparing different cellular models, we showed that ablation of METTL9 catalytic activity caused much milder neural commitment defects than full genetic abrogation of *Mettl9.* The METTL9 interactors identified in mNSCs do not contain any H[ANGST]H motifs and may form inactive complexes with the catalytic pocket of METTL9. This was confirmed by our *in vitro* methylation assays on STMN1. Taken together, these data suggest that the adequate METTL9 protein levels which preserve these protein-protein interactions might be more important than its catalytic activity, at least during early neural development. This is further documented by the observation that in *Xenopus* embryos a mutant mettl9 can rescue the neuralization phenotype with an efficiency similar to wild-type mettl9.

Moonlighting is a fascinating, emerging aspect of molecular biology. The concept is that individual proteins can possess more than one physiologically significant role. For example, in addition to their long-known and well-characterised structural role, histone octamers have also retained an evolutionarily ancient catalytic function devoted to the reduction of copper ions^117^. Moreover, several nuclear modifying enzymes also possess non-catalytic functions^118^. So far few but intriguing works have reported the existence of non-enzymatic roles for METTL proteins. For instance, N-terminus of METTL3 is sufficient to promote translation by mRNA circularization^6,119^ and METTL11A/METTL13 function as regulatory subunits of each other’s catalytic activity^120^. Recently, METTL1 was found to exert an oncogenic role in sarcomagenesis independently of its tRNA methyltransferase activity, via modulation of tRNA aminoacylation and protein synthesis^121^. Thus, we hypothesise that having both catalytic dependent and independent functions could represent a more generalised feature of METTL proteins; this could have arisen from the convergent evolution of different molecular mechanisms impinging on similar biological processes, such as the secretory pathway in the case of METTL9. Thus, our findings support the investigation of additional functions of RNA-modifying enzymes in physiologically relevant contexts.

Our protein interactome study in mNSCs showed that METTL9 binds to STMN1 and many RABs proteins, amongst the top-ranking hits. STMN1 belongs to the Stathmin family of tubulin-binding proteins and is highly expressed in the nervous system^76,77,122^. Perturbation of STMN1 impairs dendrite growth and axonal arborisation^123,124^ and is involved in regulating synaptic plasticity, memory^125^ and behaviour^126^; moreover, STMN1 levels are altered in neurodegenerative diseases and its gene is linked to a variety of psychiatric disorders^76,127^. Amongst the other METTL9 interactors, we found RAB1, RAB2, RAB7 and RAB11, which are all important for transport and secretory-related functions in neurons^128–131^; most of them are also associated with neurodegenerative or neurodevelopmental disorders^132,133^. Importantly, altered expression of STMN1, RAB1 and RAB2 has been shown to cause Golgi fragmentation^81–83,97^, similarly to METTL9 loss of function. Neural induction, axonal elongation, cell projection growth and the establishment of somatodendritic compartments heavily rely on Golgi functionality, and these processes were recurrently altered upon METTL9 down-regulation. Interestingly, alteration of Golgi morphology is often linked to impaired ER to Golgi trafficking^83,134,135^, which might explain the mis-regulation of the ER protein levels in the proteome of METTL9 deficient NSCs as well as their defective trafficking.

Proteins travelling along the secretory pathway are modified by post-translational modifications, e.g. glycosylation, whose apposition requires the precise spatio-temporal control of specific subsets of enzymes within each sub-compartment (e.g. specific Golgi cisternae)^136^. Both an intact Golgi structure and functional cellular trafficking underpin this complex process^137^. Thus, the Golgi fragmentation and altered trafficking kinetics of Golgi enzymes like ManII that we observed upon METTL9 loss could result in an aberrantly modified proteome output (mis-glycosylation)^136,137^, with a negative impact on the function of plasma membrane or secreted proteins, involved in cell-cell communication^138,139^. The delay in the ER to Golgi trafficking kinetics in *Mettl9^KO^* NSCs could also cause transient protein accumulation in the ER with negative consequences on the ER homeostasis, at the level of protein misfolding and altered protein modifications. Other parts of the secretory pathway as well as the retrograde trafficking, which are also key for recycling enzymes and sorting them correctly in each cisterna, could be affected in *Mettl9^KO^*NSCs^136,137^. Future work will be required to assess in detail the impact of METTL9 loss on cellular trafficking, by testing other RUSH cargos in METTL9 deficient mNSCs.

Our working model proposes that METTL9 sustains neural differentiation, mainly in a catalytic-independent manner, by safeguarding the secretory pathway. Mechanistically, this might occur through protein-protein interactions: upon binding to key secretory pathway regulators, like STMN1 and RABs, METTL9 might modulate them by competing with their partners and/or allosterically modifying their structure. METTL9 MTase activity has a smaller contribution to neural development, but amongst its substrates there are several Golgi-associated proteins. We showed that STMN1 outcompetes known METTL9 substrates *in vitro* and we know that STMN1 is particularly highly expressed in the developing brain^76,78^; thus, as neural development progresses *in vivo*, METTL9-STMN1 interaction could prevail over METTL9 MTase activity.

Stemming from these observations, further investigation will clarify the pleiotropic molecular roles of METTL9 in the secretory pathway, particularly during neural development.

Over 40% of the Golgi genes mutated in monogenic disorders, causing the so-called *Golgipathies*, affect the nervous system, a very high proportion compared to other tissues, and they often impair neurodevelopment^140^. Furthermore, Golgi fragmentation has been described as one of the first cellular alterations present in the neurons of patients at early stages of neurodegenerative conditions such as Parkinson’s, Amyotrophic Lateral Sclerosis and Alzheimer’s diseases, among others^141,142^, although it is not clear the extent of its contribution to the development of these diseases.

Interestingly, METTL9 expression in the adult brain peaks in the striatum, which is responsible for some of the cognitive and behavioural functions impaired in neurodegenerative diseases.

Importantly, we have highlighted a putative link between heterozygous human *METTL9* deletions and the impairment of neurodevelopment or cognitive functions in 6 patients; nevertheless, further investigation is needed to directly assess the causal relationship between *METTL9* gene loss and the onset of these neurodevelopmental disorders’ symptoms.

In conclusion, elucidating the relationship between METTL9 and Golgi function in mammalian neurogenesis will help understanding the cellular and molecular mechanisms underlying Golgi integrity and homeostasis; this, in turn, might be leveraged to amend Golgi fragmentation and dysfunction and represent an important step forward towards the treatment of both neurodevelopmental and neurodegenerative diseases.

### Data availability

Raw RNA sequencing data from mESCs and *X. laevis* experiments were submitted to the SRA archive with project identifiers PRJNA1111296 and PRJNA1111433, respectively.

Long-read whole genome sequences of the parental cell line and Mettl^KO^ clones #88 and #90 are deposited under SRA project PRJNA1242812.

The mass spectrometry proteomics data (both raw and protein groups tables) have been deposited to the ProteomeXchange Consortium^187^ via the PRIDE^188^ partner repository with the dataset identifier PXD053437.

## Acknowledgments

We thank the members of the IIT Genomics facility (Diego Vozzi, Yeraldin Chiquinquira Castillo De Spelorzi and Edoardo Henzen), the HPC Team (Sergio Decherchi, Alessandro Parodi and Mattia Pini) and the imaging facility (Michele Oneto and Marco Scotto) for their support to the research activities of the project. We are indebted to all the members of the RNA Initiative@IIT community for nurturing a stimulating scientific environment.

The pyCAG-Hygromycin plasmid was kindly gifted by Ian Chambers’ Lab, in Edinburgh. We greatly thank Professor M. Andreazzoli, E. Ferraro and M. Ori for advice and reagents for some of the procedures. This study makes use of data generated by the DECIPHER community. We are very grateful to Dr Julia Foreman, Dr Rachel Irving, Dr Vani Jain, Dr Katherine Neas, Dr Soo-Mi Park, Dr Alison Ross and Dr Vinod Varghese for kindly providing feedback on the clinical data. A full list of centres that contributed to the generation of the data is available from deciphergenomics.org/about/stats and via email from. DECIPHER is hosted by EMBL-EBI and funding for the DECIPHER project was provided by the Wellcome Trust [grant number WT223718/Z/21/Z]. The project was supported with intramural IIT funding (A. Codino, L.S., S. Gustincich and L.P.). IB and SLL were funded by the Cancer Research UK (grant reference RG86786) and by the Joseph Mitchell Fund. E.C. and F.C. were supported by the PRIN grant #2022M95RC7 from the Italian Ministry of University and Research (MUR) and the Tuscany Health Ecosystem – THE grant from MUR.

## Author contributions

A. Codino designed, performed and analysed the experiments and wrote the manuscript. L.S. contributed to the execution of neural differentiation protocols, immunofluorescence and molecular cloning experiments. C.O. performed the manipulations of *Xenopus laevis* embryos, with the help and under the supervision of R.V. A. Cuomo performed the mass spectrometry analysis for proteomics experiments. H.S.R performed the *in vitro* methyltransferase assays. M.P. and N.M. performed the acidic hydrolysis experiments for the detection of 1MH, 3MH and His in NSCs extracts, under the supervision of S.Girotto and R.S. S.L performed the experiments for the generation of the *Mettl9^KO^* mESC line under the supervision of I.B. E.C. performed some of the neural differentiation experiments under the supervision of F.C. P.B. supervised the microscopy analysis. R.R. co-supervised A.C. in the RUSH and digitonin-IF experiments. A.B., S. Gustincich and T.K. provided help and support for the molecular biology techniques of the project. L.P. conceived and supervised the study, performed the bioinformatics analysis and wrote the manuscript. All the authors revised and approved the manuscript.

**Supplementary Fig. 1:**
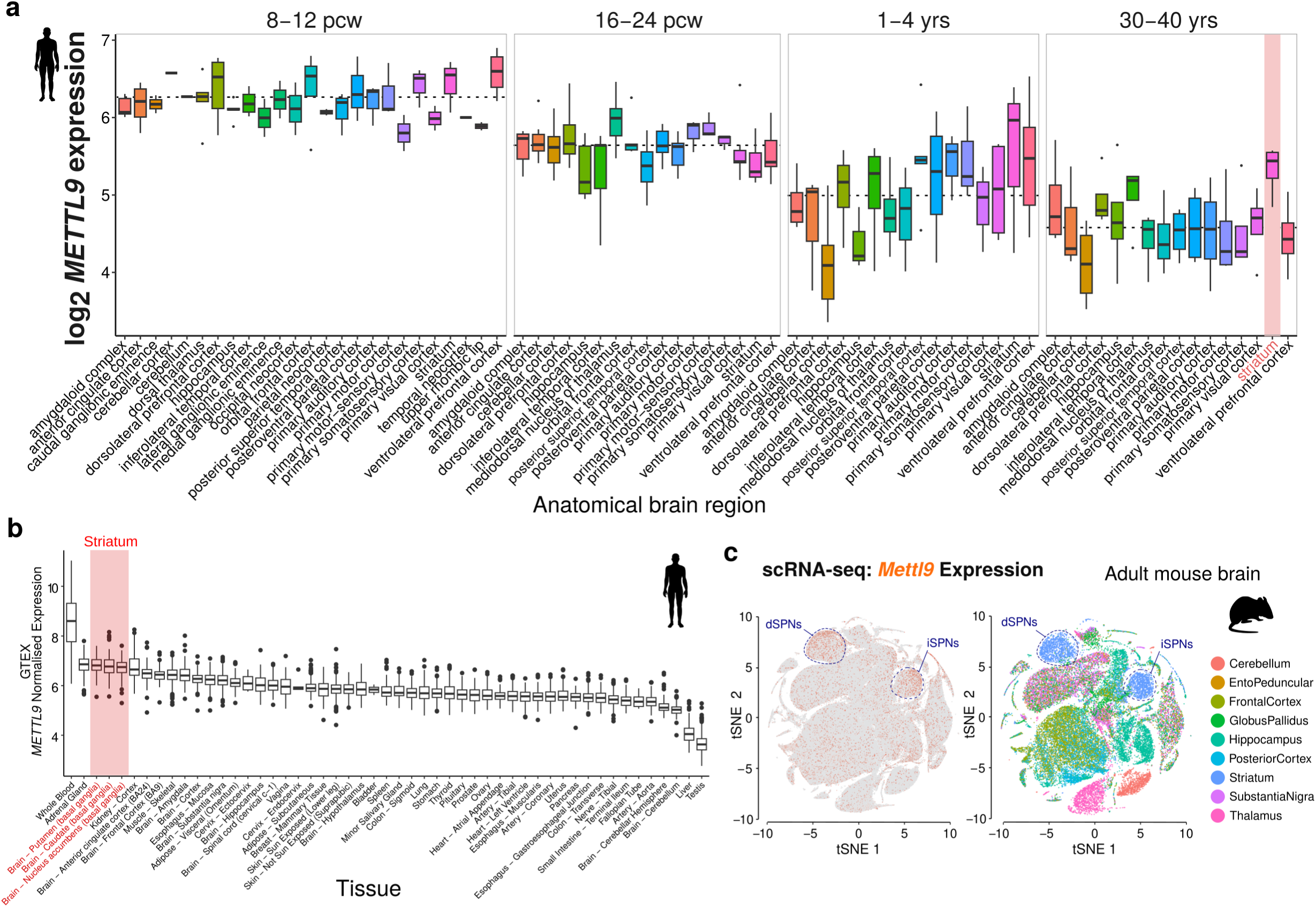
Mettl9 expression in human brain development and mouse adult brain. **a** *METTL9* expression (log_2_) in pre-natal (8-12 and 16-24 post conception weeks), and post-natal (1-4 years: yrs, 30-40 yrs) brain regions. The pink selection highlights the anatomical area with highest *METTL9* expression in adulthood (BrainSpan Atlas of the Developing Human Brain). **b** Normalised *METTL9* expression within RNA-seq datasets from human tissues (GTEx); the striatum (putamen, caudate, nucleus accumbens) is highlighted in pink. **c** Re-analysis of scRNA-seq data (Saunders et al. 2018) of adult mouse brain; on the left, tSNE plot with *Mettl9* expression (in orange) in direct and indirect spiny projection neurons (dSPNs and iSPNs).

**Supplementary Fig. 2:**
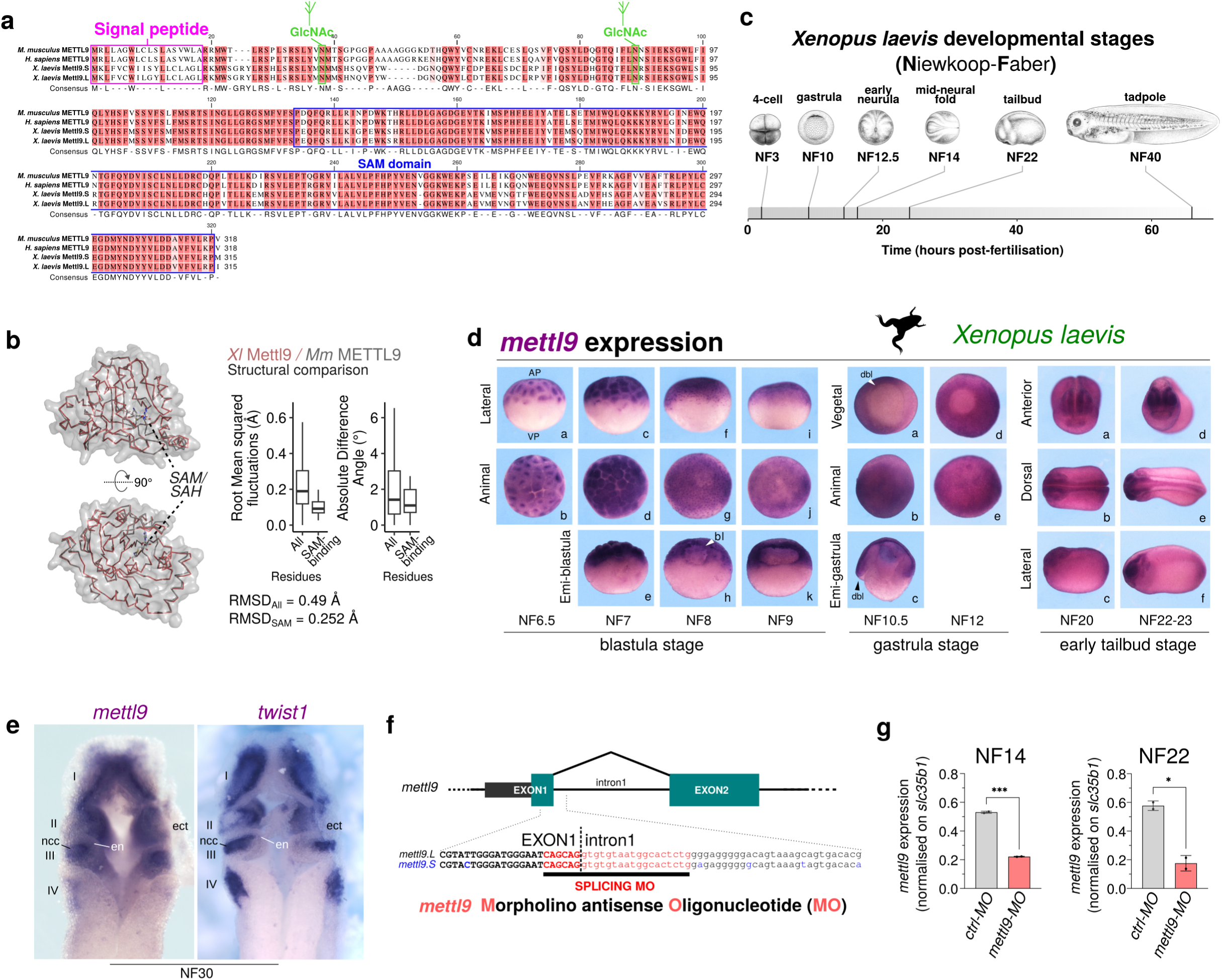
*mettl9* expression and knock down strategy in *X. laevis*. **a** Protein alignment of mouse, human, and *X. laevis* S and L (Short and Long chromosome) METTL9/Mettl9. The signal peptide predicted site, the predicted N-Acetylglucosamine (GlcNAc) deposition sites, and the SAM domain are highlighted. **b** Structural comparison of *X. laevis* (*Xl*) Mettl9 and *M. musculus (Mm)* METTL9 proteins (the backbones of the AlphaFold predictions are superimposed in red and gray, respectively). Boxplots on the right show the distributions of root mean squared fluctuations and pseudo-torsion angle deviations for all residues (All) and amino acids engaged in SAM binding (SAM-binding, including D151/G153). Global root mean squared deviations (RMSD) are indicated in the lower part. **c** Main developmental stages of *X. laevis* (adapted from (Zahn et al. 2022)). **d** *mettl9* mRNA expression at blastula (NF 6.5-9), gastrula (10.5-12) and early tailbud (NF20-23) stages, shown by RNA WISH. AP: Animal Pole; VP: Vegetal Pole; bl: blastocoel; dbl: dorsal blastopore lip. **e** *mettl9* and *twist1* expression at stage NF30 in horizontally sectioned embryos, shown by WISH. The core of the pharyngeal arches (I-IV) (neural crest cells, ncc), endoderm (en) and ectoderm (ect) are shown. **f** *mettl9* Morpholino antisense Oligonucleotide (MO) targeting the exon1-intron1 junction of *X. laevis mettl9* pre-mRNA. **g** Relative expression of *mettl9* mRNA (by qPCR) normalised on *β-actin*, at NF14 and NF22 stages, in control (*ctrl*-) and *mettl9*-MO embryos. (t-test).

**Supplementary Fig. 3:**
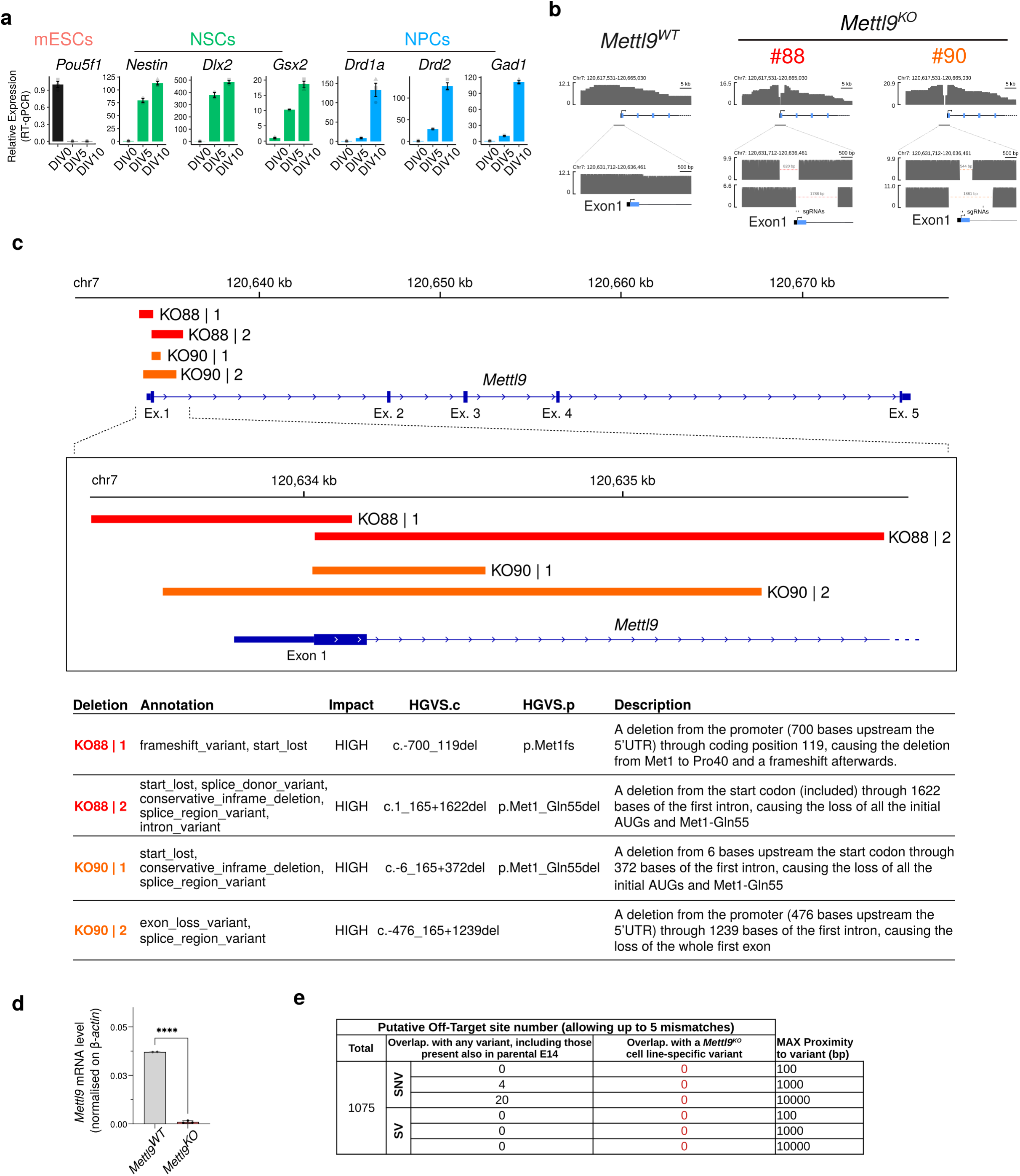
Validation of the *Mettl9^KO^* mESC lines. **a** Relative mRNA expression level (by qPCR) of mESC, NSCs or NPCs marker genes at DIV0, DIV5 and DIV10 during mESC neural differentiation. **b** Validation of the *Mettl9* (Exon1) deletions by long-read sequencing, in the *Mettl9^KO^* mESC lines (#88 and #90), generated by CRISPR/Cas9. **c** Genomic diagram showing the overall *Mettl9* gene structure (top) and a close-up (bottom) of the exact positions of the biallelic deletions in *Mettl9^KO^* #88 and #90 cell lines (red and orange, respectively). The lower table provides a detailed annotation of the functional consequences due to each deletion, including their nomenclature, which follows the HGVS (Human Genome Variation Society) standard. **d** Relative *Mettl9* mRNA expression level (qPCR) in *Mettl9^WT^* and *Mettl9^KO^*NPCs. (t-test). **e** Table summarising the total number of putative off-target (OT) sites (OT; left column), and the number of OT overlapping either ubiquitous (center column) or *Mettl9^KO^*-specific variants (SNV: single-nucleotide variants; SV: structural variants). These figures are calculated taking into consideration a 100, 1000 and 10000-bp window around each putative OT.

**Supplementary Fig. 4:**
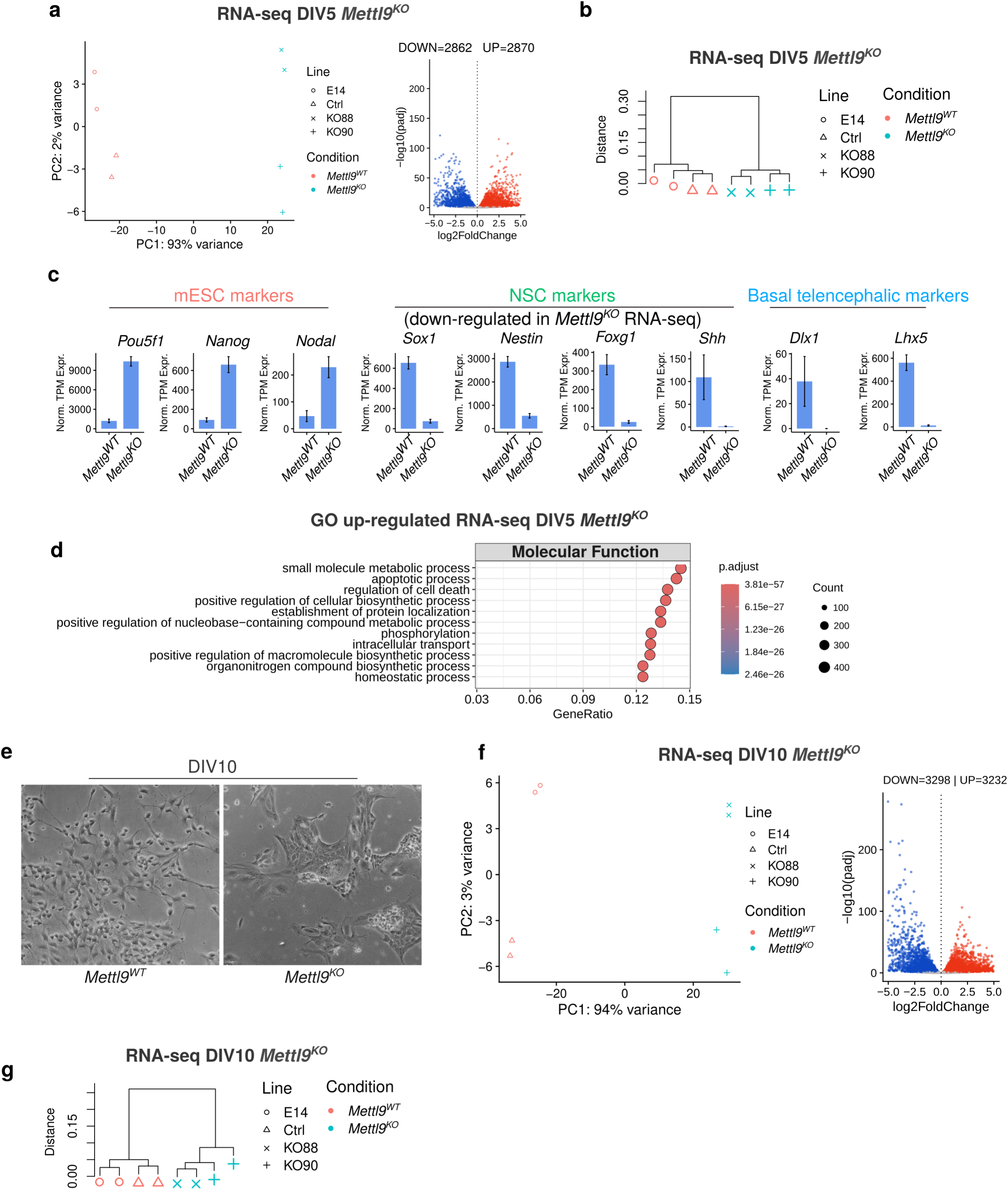
Phenotypic characterisation of the *Mettl9^KO^*mESC lines at DIV5 and DIV10. **a** Principal component analysis (PCA) plot (left) and volcano plot (right) of Differentially Expressed Genes (DEG) in *Mettl9^WT^* and *Mettl9^KO^* NSCs (DIV5 RNA-seq). **b** Hierarchical clustering analysis showing very high consistency in global gene expression values between *Mettl9^WT^* cells (i.e. the parental E14 and a non-edited clonal control line), and between *Mettl9^KO^*clones (i.e. #88 and #90) at DIV5, as indicated by the low similarity distance values within each of these groups. **c** Normalised TPM expression of mESC (light red), NSCs (green) and NPCs (light blue) marker genes. **d** Top (10) GO Molecular Function terms up-regulated in *Mettl9^KO^* RNA-seq (DIV10). **e** Representative brightfield images of *Mettl9^WT^* and *Mettl9^KO^* NPCs at DIV10. **f** PCA plot (left) and volcano plot (right) of Differentially Expressed Genes (DEG) in *Mettl9^WT^* and *Mettl9^KO^* NPCs (RNA-seq DIV10). **g** Hierarchical clustering analysis for cells at DIV10, following the same criteria as in panel **b** (DIV5), confirming a high consistency within *Mettl9^WT^* and *Mettl9^KO^* cell groups also at a later timepoint of differentiation.

**Supplementary Fig. 5:**
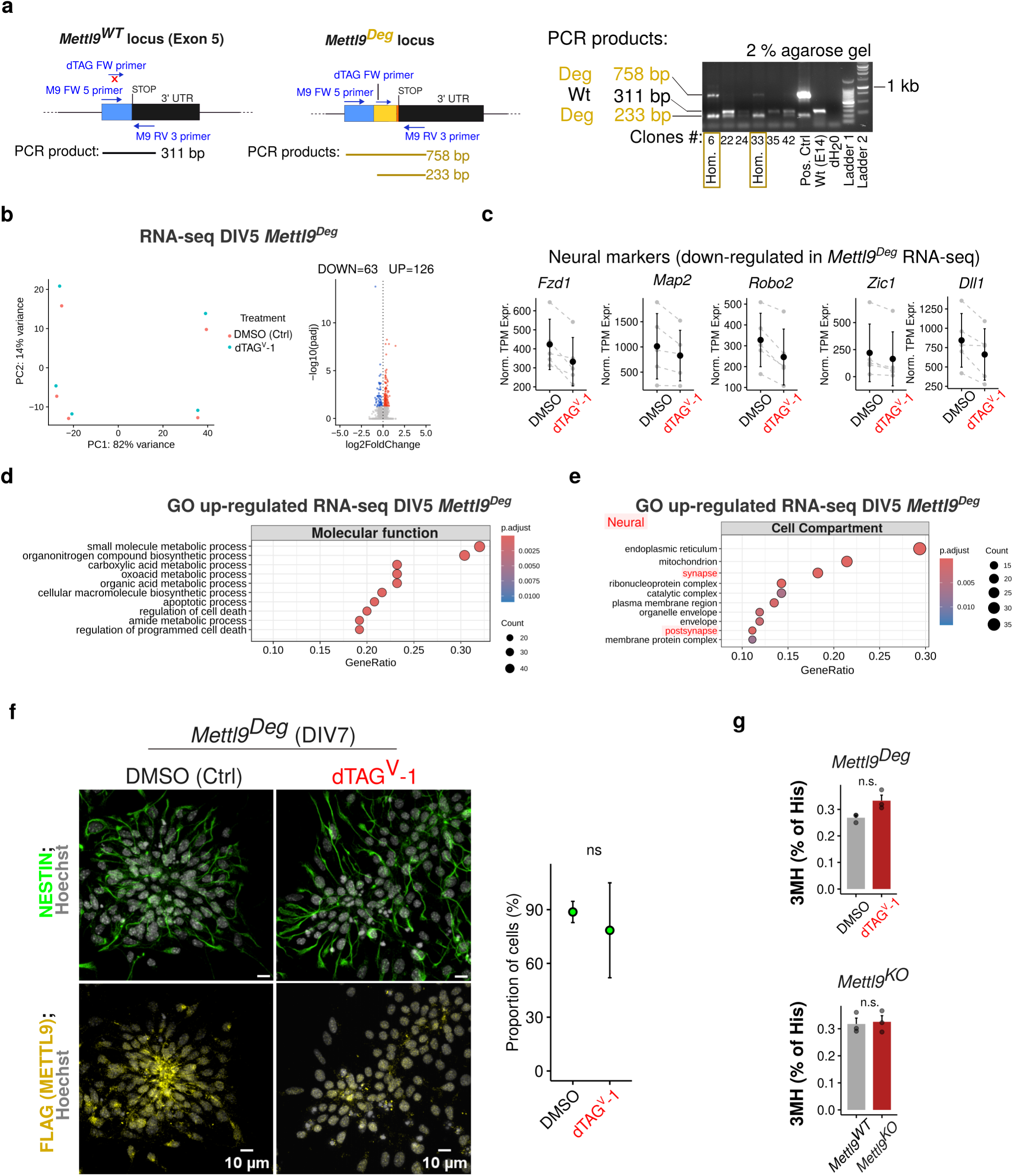
Characterization and transcriptomic analysis of the *Mettl9^Deg^* mESC line. **a** Genotyping strategy used to screen *Mettl9^Deg^*mESC clones. On the left, the schematic depicts the amplification of the 311 bp PCR product from the *Mettl9^WT^* locus (where only 2 primers out of three anneal); the central part of the panel shows that PCR amplification of the *Mettl9^Deg^*locus generates 2 products of 758 bp and 233 bp. On the right, an agarose gel shows examples of PCR products from homozygous (Hom.) *Mettl9^Deg^* ESC clones (#6 and #33, rectangles) with 2 bands (the highest at 758 bp and lowest at 233 bp). The positive control (Pos. Ctrl) is the targeting vector; PCR-amplification of untargeted *Mettl9^WT^* locus generates only one band (311 bp), as shown in the WT (E14) lane. DNA ladder 1 and 2 are respectively 100 bp and 1 kb ladders. ***b*** PCA (left) and volcano plot (right) of Differentially Expressed Genes (DEG) in *Mettl9^Deg^* NPCs (DIV5 RNA-seq). **c** Normalised TPM expression of neural marker genes from *Mettl9^Deg^*RNA-seq (DIV5). **d** Top 10 up-regulated GO Molecular function terms in *Mettl9^Deg^* (RNA-seq, DIV5). **e** Top up-regulated Cellular Component GO terms in *Mettl9^Deg^* RNA-seq experiment (RNA-seq, DIV5). **f** Representative immunofluorescence (IF) images of DMSO– and dTAG^V^-1 treated *Mettl9^Deg^* NSCs (DIV7), stained with an anti-NESTIN antibody (green), Hoechst (grey), and anti-FLAG antibody (yellow). Scale bar is 10 μm; relative quantification of NESTIN signal on the right (Number of cells counted: 2697 for *Mettl9^WT^*; 1839 for *Mettl9^KO^*. Wilcoxon test). **g** Relative bulk 3MH levels (% of total histidine) quantified by mass spectrometry in *Mettl9^KO^* and dTAG^V^-1 treated *Mettl9^Deg^* compared to *Mettl9^WT^* and DMSO-treated *Mettl9^Deg^* (respectively) NSCs at DIV6. (Wilcoxon test).

**Supplementary Fig. 6:**
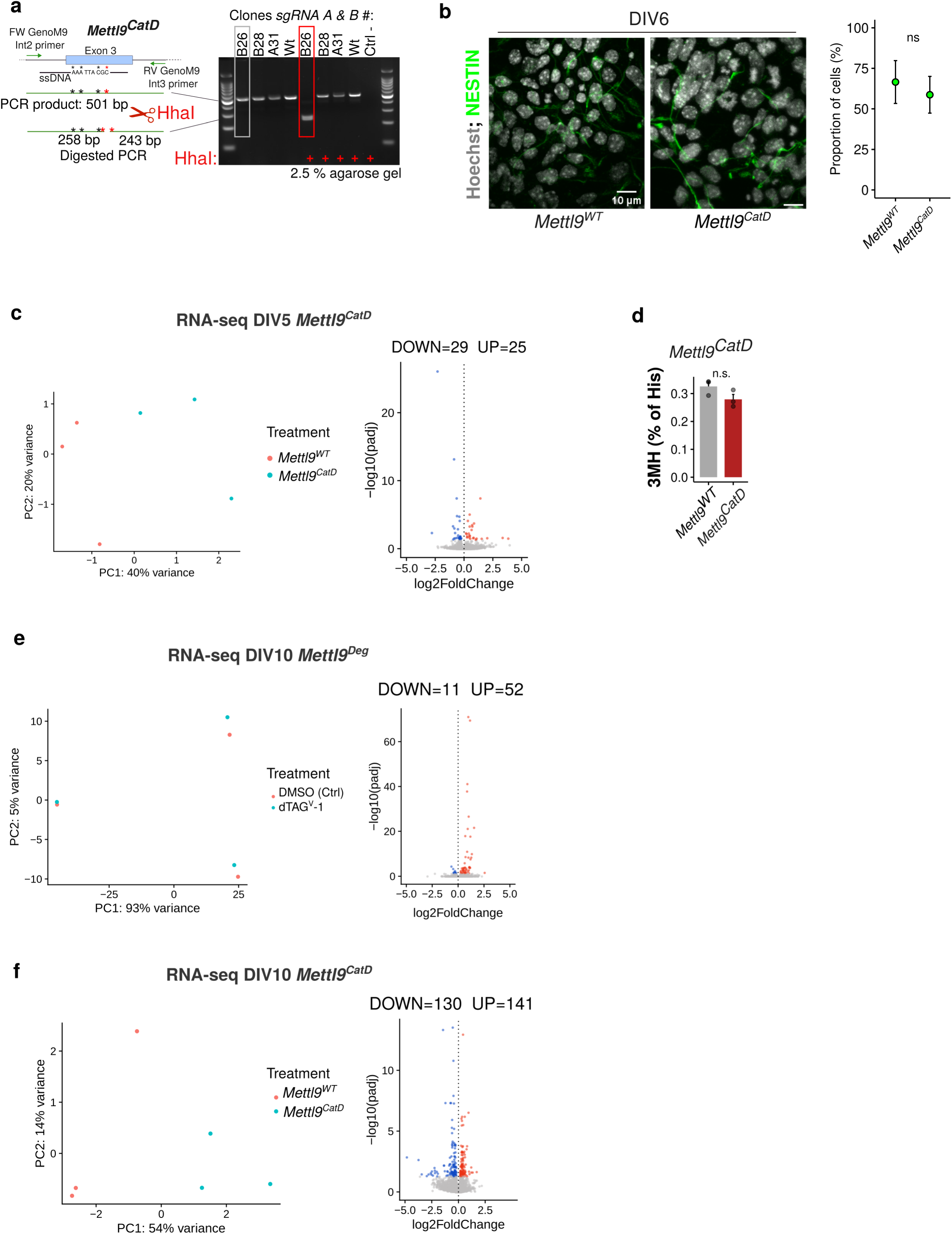
Characterisation of the *Mettl9^CatD^* mESC line and of METTL9 catalytic dependent roles in neural development. **a** Genotyping strategy adopted to screen *Mettl9^CatD^* mESC clones. On the left, the schematic depicts the PCR amplification of the targeted *Mettl9^CatD^* locus (asterisks are the 4 desired mutations), which generate a 501 bp product. Upon HhaI addition, only mutated PCR products (harboring the fourth mutation, red asterisk) are cut and generate 2 DNA fragments of 258 bp and 243 bp. On the right, an agarose gel shows the 501 bp PCR product from a homozygous *Mettl9^CatD^* clone (#B26, rectangle), which is shifted after HhaI digestion (double band, 258 and 243 bp). HhaI-resistant PCR products identify DNA from *Mettl9^WT^* clones (e.g. #B28). **b** Representative IF images of *Mettl9^CatD^* DIV5 NSCs stained with anti-NESTIN antibody and Hoechst. Scale bar is 10 μm; relative quantification of NESTIN signal on the right (Number of cells counted: 1647 for *Mettl9^WT^*; 1645 for *Mettl9^CatD^*. Wilcoxon test). **c** PCA (left) and volcano plot (right) of differentially expressed genes in *Mettl9^WT^* and *Mettl9^CatD^* NSCs (DIV5 RNA-seq). **d** Relative 3MH levels (% of histidines) in *Mettl9^WT^* and *Mettl9^CatD^* NSCs quantified by mass spectrometry. **e** PCA (left) and volcano plot (right) of differentially expressed genes in Ctrl (DMSO-) and dTAG^V^-1-treated *Mettl9^Deg^* NPCs (DIV10 RNA-seq). **f** PCA (left) and volcano plot (right) of differentially expressed genes in *Mettl9^WT^*and *Mettl9^CatD^* NPCs (DIV10 RNA-seq).

**Supplementary Fig. 7:**
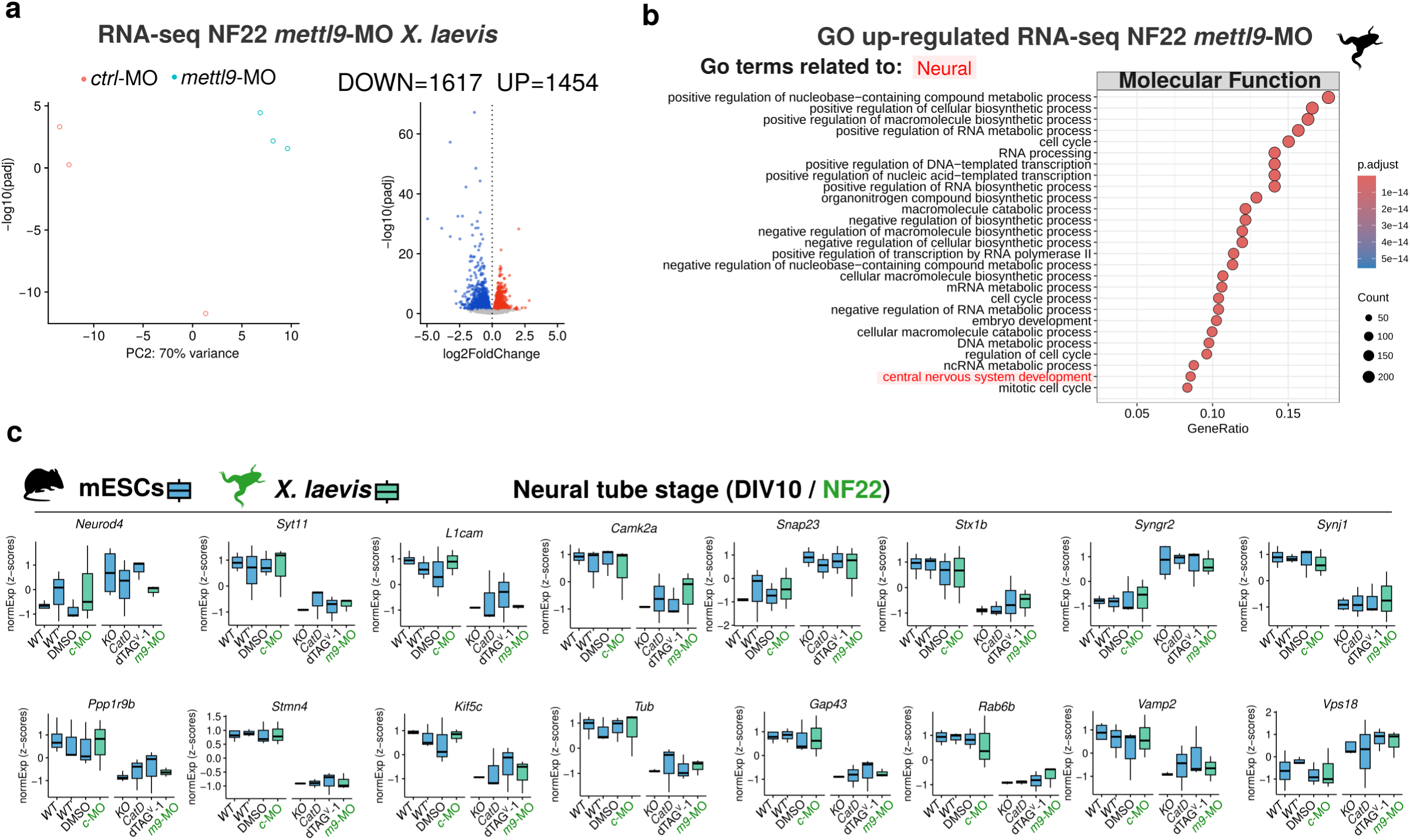
Transcriptomic characterisation *X. laevis mettl9*-MO embryos at NF22 stage and comparison with the mouse NPCs lines. **a** PCA (left) and volcano plot (right) of differentially expressed genes in *mettl9*-MO embryos and *ctrl*-MO (NF22 RNA-seq). **b** Top up-regulated Molecular Function GO terms in *mettl9*-MO NF22 embryos. **c** Differentially expressed genes involved in neural processes consistently mis-regulated at DIV10 (mESCs) and NF22 (*X. laevis*). *WT’* is the clonal WT control for *Mettl9^CatD^*. *c-*MO and *m9*-MO are *ctrl*-MO and *mettl9*-MO, respectively.

**Supplementary Fig. 8:**
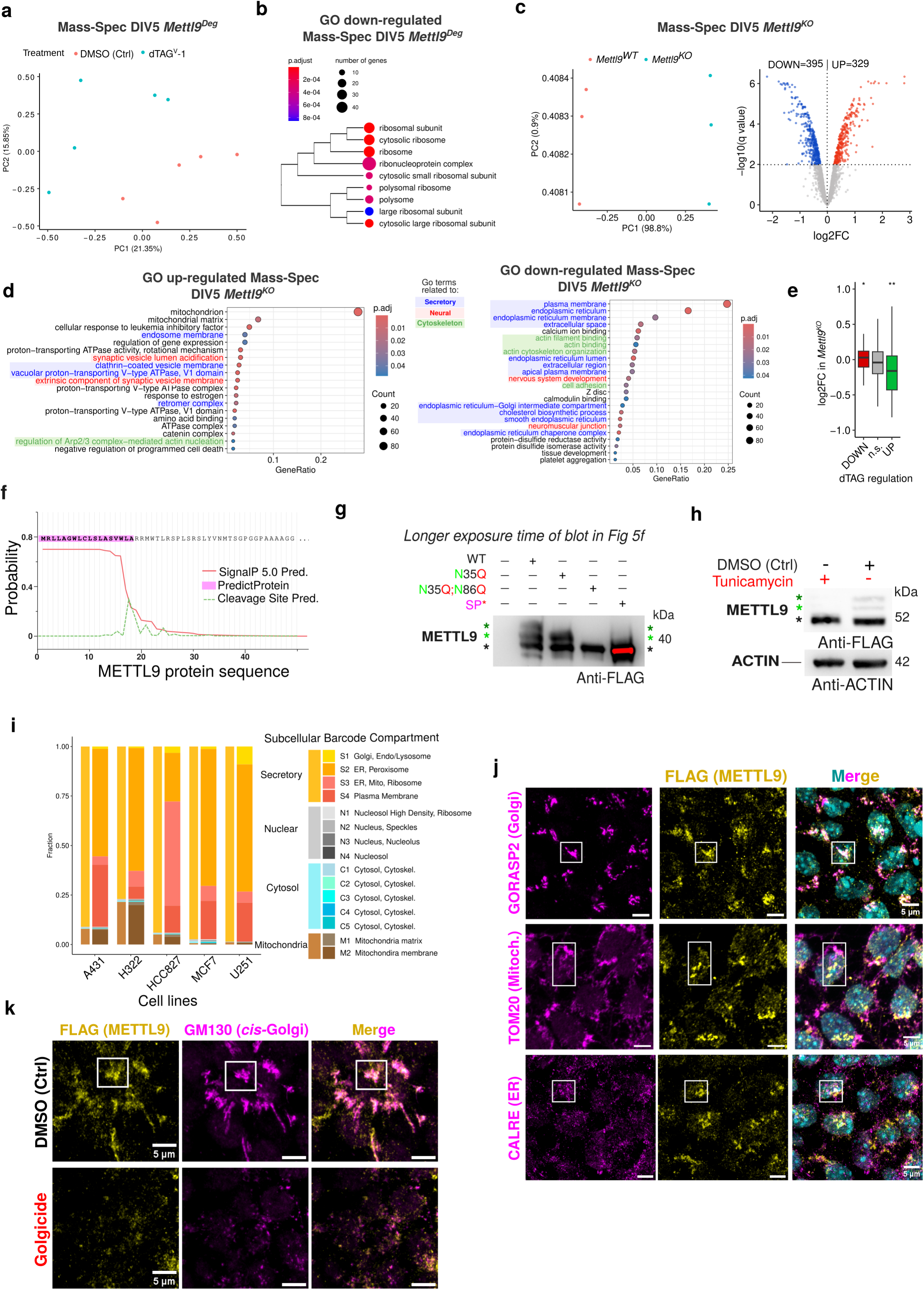
Proteomic analysis upon acute or constitutive METTL9 depletion in NSCs and METTL9 subcellular localization in NSCs. **a,b** PCA (a) and GO analysis (**a**) on proteomic dataset of acutely depleted METTL9-DEG in *Mettl9^Deg^* NSCs. GO terms (**b**) refer to the down-regulated proteins. **c-d** Volcano plot (**c**) and GO analysis (**d**) of the misregulated proteins (coloured dots) identified by mass spectrometry in *Mettl9^KO^* NSCs over *Mettl9^WT^* (ctrl). Up-regulated genes with an adjusted p value < 0.2 were used for this analysis. **e** Comparison between the misregulated proteins of dTAG^V^-1-treated *Mettl9^Deg^* NSCs and *Mettl9^KO^* (* p<0.05; ** p<0.01; Wilcoxon test). **f** Signal peptide and cleavage sites prediction within METTL9 amino acid sequence, with Signal IP 5.0 and PredictProtein (Almagro Armenteros et al. 2019; Bernhofer et al. 2021). **g** Longer exposure of the same WB membrane (Anti-FLAG) as displayed in Fig. 5f highlighting the absence of higher METTL9 bands (green asterisks) in the 3 mutated METTL9 (METTL9-N35Q-FLAG (N35Q), METTL9-N35Q;N86Q-FLAG (N35Q;N86Q), and SP*-METTL9-FLAG (SP*). An Anti-FLAG antibody was used. Black, green and dark green asterisks (*) refer to the lowest, intermediate and top METTL9 bands, respectively. **h** WB showing METTL9-DEG (visualised with an anti-FLAG antibody) in tunicamycin-treated mESCs extracts. An anti-ACTIN antibody was used as a loading control. Asterisks (*) as in (**g**). **i** Relative proportion of METTL9 found in distinct subcellular fractions (each colour refers to a different compartment) in proteomic data from (Orre et al. 2019). **j** Representative IF images of *Mettl9^Deg^* NSCs co-stained with anti-FLAG antibody for METTL9 and: anti-GORASP2 (Golgi) or anti-TOM20 (mitochondria) or anti-CALRE (ER). Nuclei are stained by Hoechst (cyan). Scale bar is 5 μm. **k** Representative IF images of *Mettl9^Deg^* DMSO-or golgicide-treated NSCs (DIV6), co-stained with anti-FLAG antibody (METTL9) and anti-GM130 (Golgi). Scale bar is 5 μm.

**Supplementary Fig. 9:**
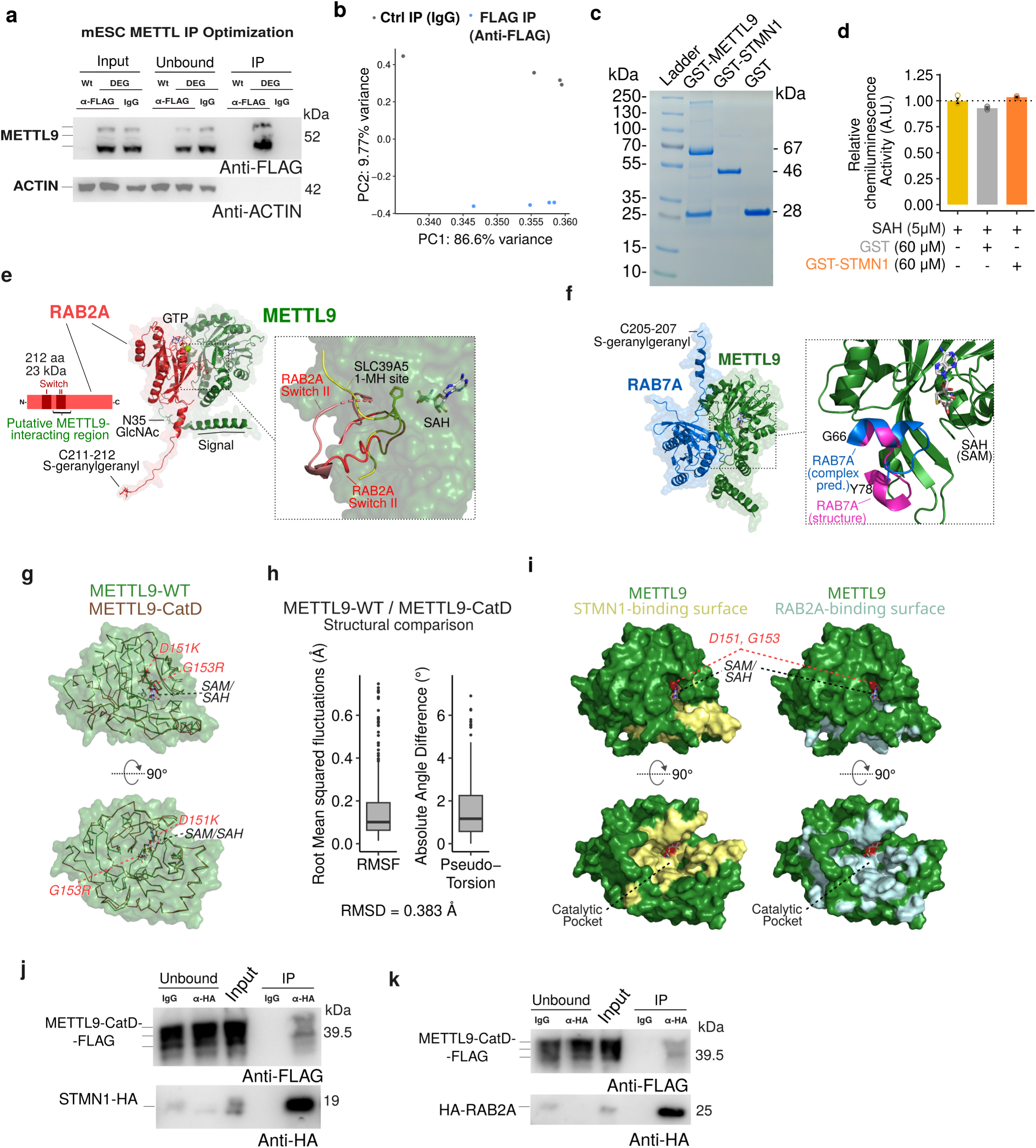
Characterisation of METTL9 protein interactors in mNSCs. **a** WB showing a representative METTL9 immunoprecipitation (IP) with anti-FLAG antibody or anti-IgG (Ctrl), from *Mettl9^WT^* (WT) and *Mettl9^Deg^* (DEG) mESCs. **b** PCA of METTL9 IP-MS samples at DIV4 (NSCs). **c** Coomassie-stained poly-acrylamide gel showing the recombinant GST-METTL9 (GST-METTL9-FLAG), GST-STMN1 (GST-STMN1-HA) and GST proteins after affinity purification from *E. coli* extracts. **d** *In vitro* control of the detection method used in methyltransferase assays, showing that the recombinant GST-STMN1 or GST proteins do not interfere with the downstream conversion of S-adenosyl-homocysteine (SAH) into chemiluminescence activity. Concentrations (μM) of each component are shown. **e**, **f** AlphaFold modelling prediction of RAB2A-METTL9 (**e**) or RAB7-METTL9 (**f**) protein complexes. (RAB2A in red; RAB7 in blue; METTL9 in green). **g** Superimposed structures of METTL9-WT (green) and METTL9-CatD (brown backbone), as predicted by AlphaFold. **h** Comparison of the METTL9-WT and METTL9-CatD structures depicted in **g**, representing the distribution of root mean squared fluctuations and pseudo-torsion angle deviations. The overall root mean square deviation value (RMSD) is shown in the lower part. **i** Representation of the relative positions of the D151;G153 residues mutated in METTL9-CatD (red) and the predicted interaction surfaces between METTL9 (green) and STMN1 (yellow) or RAB2A (light blue). **j, k** WB showing the immunoprecipitation (IP) of STMN1-HA (**j**) or HA-RAB2A (**k**) with anti-HA beads (HA) or IgG (ctrl), after co-expression of STMN1-HA (**j**) or HA-RAB2a (**k**) and METTL9-CatD-FLAG in mESCs (visualised with anti-FLAG and anti-HA antibodies).

**Supplementary Fig. 10:**
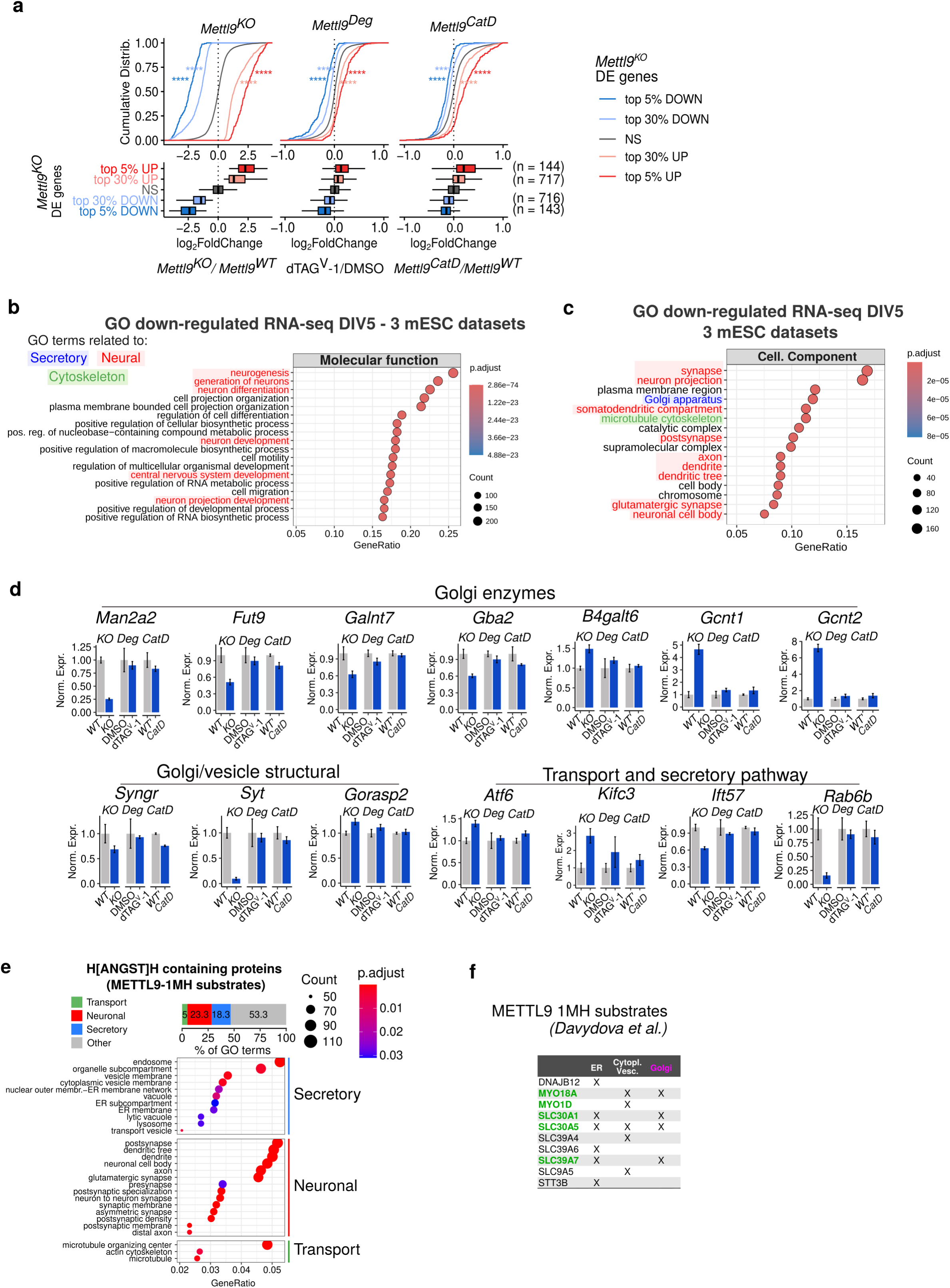
Biological processes consistently affected across *Mettl9^KO^/^CatD^/^Deg^* mNSCs and *in silico* analysis of METTL9 substrates. **a** Graphs tracking how the genes mis-regulated upon *Mettl9^KO^*(represented in the four different colors) behave in the other mESC lines under study. The distribution of the (log2) fold changes for each gene group is shown by means of both *cumulative distribution* (top) and *box* plots (bottom). This analysis reveals that up– and down-regulated genes in *Mettl9^KO^* are generally similarly affected (i.e. they display a concordant, statistically significant up-or down-regulation trend compared to control) also in *Mettl9^Deg^*and *Mettl9^CatD^*, albeit to a much smaller extent. For each of the four gene groups considered, the panel also shows its number (on the right of the corresponding boxplot line). **b-c** Most enriched Molecular Function (b) and Cellular Component (c) GO terms for the significantly down-regulated genes across the three mouse cell lines. **d** Example of a subset of differentially expressed genes encoding for Golgi enzymes; structural Golgi and vesicles’ proteins and transport/secretory pathway-related proteins in *Mettl9^KO^*, dTAG^V^-1-treated *Mettl9^Deg^* and *Mettl9^CatD^* lines and their relative controls (DIV5). **e** Some of the most enriched GO terms (Transport, Endomembrane and Neuronal) in H[ANGST]H-containing mouse proteins (i.e. potential METTL9 substrates). **f** METTL9-1MH substrates, experimentally validated by Davydova et al. (Davydova et al. 2021), whose localisation is within ER, Cytoplasmic Vesicles (i.e. endosomes, secretory/synaptic vesicles) and Golgi.

**Supplementary Fig. 11:**
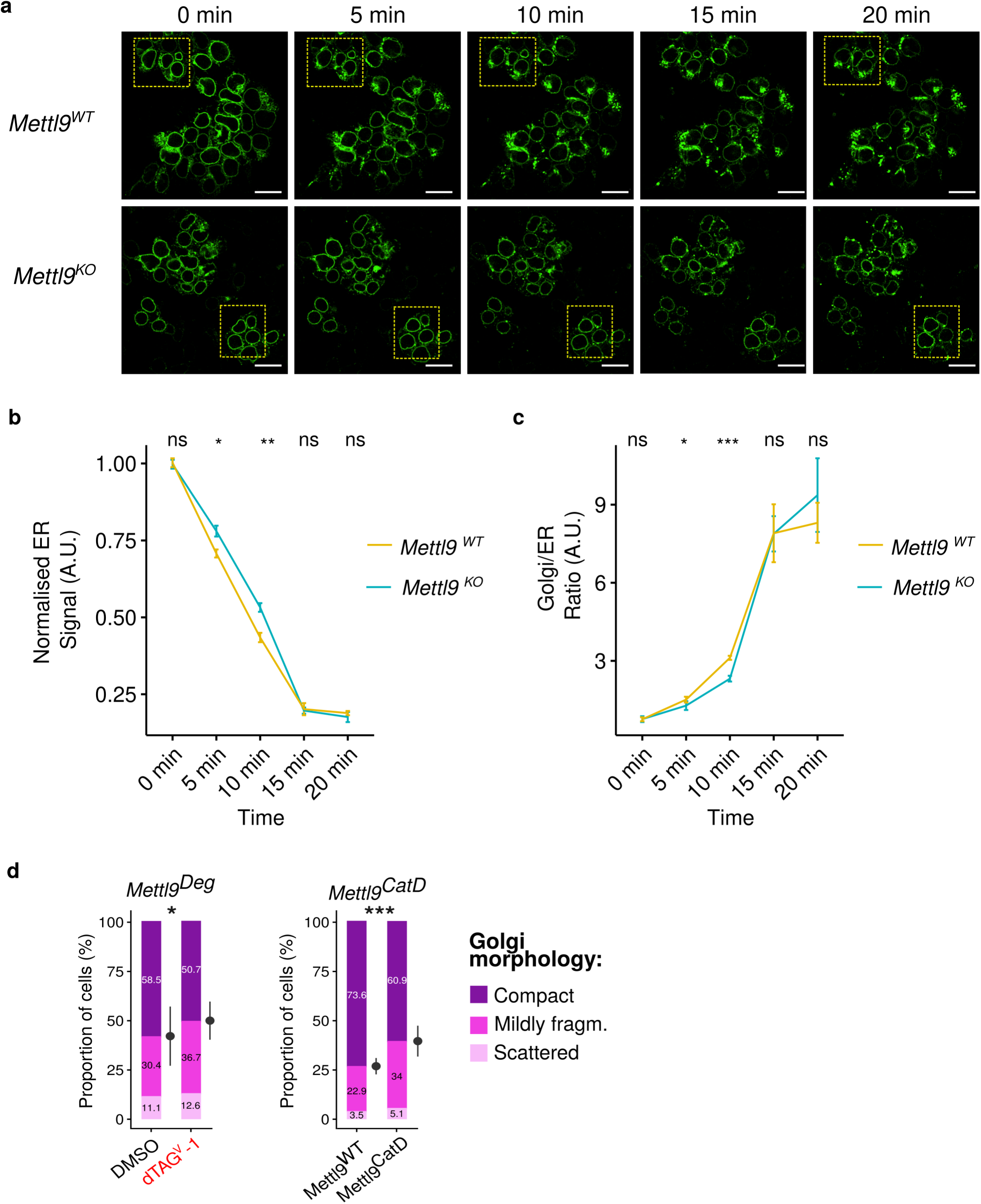
Cellular trafficking analysis by RUSH in *Mettl9^KO^* NSCs and quantification of Golgi morphology in *Mettl9^Deg^* and *Mettl9^CatD^*. **a** Entire fields of view of the corresponding zoomed-in images displayed in Fig.8**b**, showing ManII-SBP-EGFP signal in *Mettl9^WT^* and *Mettl9^KO^* NSCs, at 0, 5, 10, 15 and 20 minutes after Biotin addition. **b,c** Normalised ManII-SBP-EGFP signal in the ER (b) and Golgi over ER signal ratio (c) in *Mettl9^WT^*and *Mettl9^KO^* NSCs across 3 independent RUSH experiments. (*: P<0.05, **: P<0.01; ***: P<0.001; T test). **d** Relative number of cells displaying a compact, mildly fragmented or scattered Golgi in dTAG^V^-1-treated *Mettl9^Deg^* compared to DMSO-treated NSCs and in *Mettl9^CatD^*or *Mettl9^WT^* mNSCs. Number of cells counted: 773 (DMSO), 532 (dTAG^V^-1); 608 (*Mettl9^WT^*) and 749 (*Mettl9^CatD^*); (χ2 test).

**Supplementary Fig. 12.**
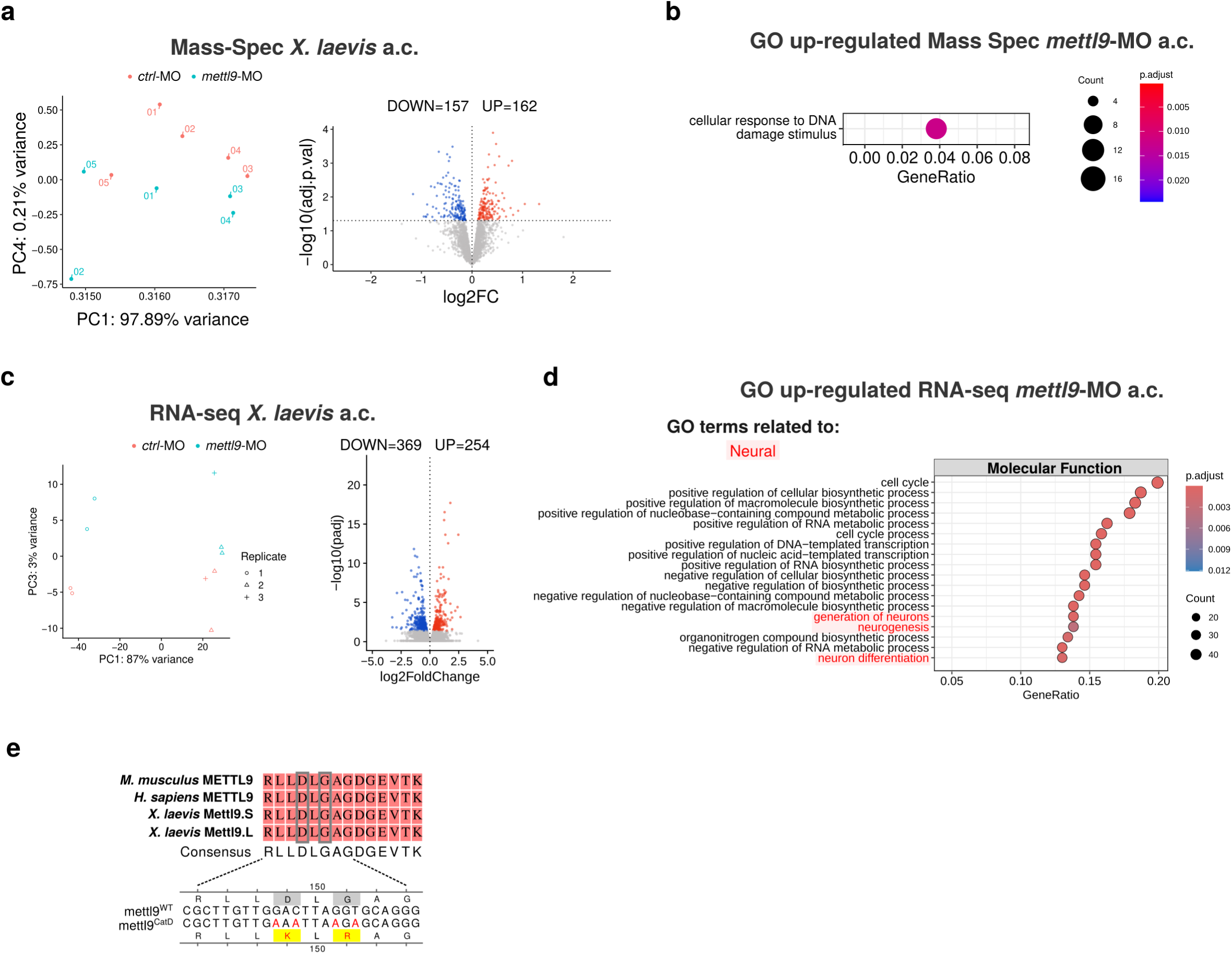
Proteomic and transcriptomic analysis of *Mettl9*-depleted *X. laevis* animal caps (a.c.) and design of *mettl9^CatD^* mRNA for the rescue. **a** PCA (left) and volcano (right) plots of mis-regulated proteins in *mettl9*-MO neuralised animal caps (a.c.) vs. *ctrl*-MO neuralised a.c. **b** Up-regulated GO term in the *mettl*-MO proteome versus *ctrl*-MO proteome of neuralised a.c. **c** PCA (left) and volcano (right) plots of differentially expressed genes in the *mettl9*-MO versus control neuralised animal caps RNA-seq experiment. **d** Most up-regulated Molecular Function GO terms of DEGs in the same experiment described in (**c**). **e** Schematic showing the 4 nucleotides mutagenised in *mettl9^CatD^* mRNA encoding D149K and G151R within the conserved amino acid sequence of *X. laevis* Mettl9 catalytic domain.

